# Intracranial Insights into the Developing Neural Basis of Moral Punishment

**DOI:** 10.1101/2025.07.27.667012

**Authors:** Keyu Hu, Haoming Zhang, Jiayu Cheng, Yangchun Chen, Xiaobin Zhang, Changlin Bai, Fengpeng Wang, Yi Yao, Haiyan Wu

## Abstract

Third-party punishment (TPP) is a critical component of social regulation and justice, integrating moral reasoning and emotional salience. However, the neural developmental basis underpinning this complex process remains largely unknown. Using rare intracranial stereo-electroencephalography (SEEG) in 14 children and 17 adults, we investigated the developing neural circuits of TPP. We found that broad-band gamma activity in the amygdala and ventromedial prefrontal cortex (vmPFC) encodes inferred intentions, with different patterns across age groups. Furthermore, the vmPFC, insula, and inferior parietal lobule (IPL) integrate punishment efficacy, also showing significant developmental differences. Combining task and resting state functional connectivity analyses, we further found age-dependent interactions among the amygdala-insula and IPL-vmPFC neural couplings during decision-making. These findings provide valuable intracranial evidence that the maturation of moral decision-making stems from the developmental refinement of subcortical-cortical circuits that integrate emotional and cognitive evaluations, explaining the shift from intuitive decisions in children to context-sensitive judgments in adults.

## 1 Introduction

As a common decision individuals face in real life, third-party punishment (TPP) refers to the act of punishing transgressors who harm others even when they are not directly harmed by the wrongdoings[1, 2, 3, 4]. TPP of misconducted behaviors has been considered a distinctive trait of human societies and is universal across cultures [5]. While studies have shown the capacity for social evaluation and a rudimentary understanding of moral principles develop very early [6, 7, 4], comprehensive TPP decision requires integrating the consideration of multiple sources of information, including the intention behind an action and the efficacy of punishment in producing desired outcomes [8, 9]. The ability to combine these elements is thought to develop progressively over time.

Among the key factors guiding punishment decisions, inferred intention plays a central role. Observers tend to assign greater moral weight to actions perceived as more intentional, even when outcomes remain the same[10, 11, 12, 13]. This bias toward intentional harm is well documented in both first-person and third-party contexts[14, 15, 16, 17], which underscores intention as a core determinant of moral judgment and punishment. Beyond inferred intentions, the consequences or outcomes of an action also critically influence moral decisions [18]. Specifically, within the TPP context, the efficacy of the punishment, which refers to the perceived effectiveness of the punishment in achieving the desired outcome, can also influence the decision of the punishers. Previous research shows that individuals are more likely to support punitive measures when they believe these measures will effectively prevent future harm [19, 18]. Further, people attribute the importance of outcomes to automatic emotional processes, suggesting that one’s moral decisions are deeply influenced by the emotional responses to the consequences of action [20].

At the neural level, previous studies have shown that TPP decisions require the collaborative effort of multiple systems involved in affective, evaluative, and social cognitive processing[21, 22, 16]. From the affective perspective, TPP is associated with emotion brain especially brain regions involving the negative emotion processing [23, 20, 9]. Studies consistently indicate that the insula and amygdala are key regions involved in the generation and processing of emotional experiences that guide behavior[21, 24, 25, 26]. Meanwhile, empathy towards victims is also an important component to drive punitive actions, and the amygdala and insula are also considered to play important roles in the human empathy network[27, 28, 29]. These affective responses may then be integrated and evaluated in the ventromedial prefrontal cortex (vmPFC), which is essential for value integration and decision-making [9, 30, 31, 32, 33, 17]. Additionally, the inferior parietal lobule (IPL) has been consistently implicated in theory of mind (ToM) and moral judgments, highlighting its contribution to the multifaceted nature of moral decision-making [15, 34]. Furthermore, the IPL is recognized as a marker for expectation violation and memory retrieval, both crucial processes in social information processing [35].

Beyond local regional involvement, a critical aspect of complex cognitive processes like TPP lies not just in the activation of specific areas, but also in the dynamic reconfiguration of their functional relationships. While the brain maintains a stable, intrinsic functional architecture during resting states, serving as a crucial baseline [36], neural circuits may undergo significant re-organization during demanding tasks, diverging from this baseline to meet novel processing demands [37, 38]. Quantifying the extent and nature of these task-induced reconfigurations, relative to the intrinsic resting state, offers deeper mechanistic insights into how the brain dynamically adapts to complex social computations.

Despite growing insights from conventional neuroimaging studies, two critical questions remain unresolved. First, how do neural representations of intention and efficacy develop across childhood and adolescence in the context of TPP? Second, how does inter-regional connectivity reorganize over development to support the shift from intuitive, emotion-driven responses to more integrative, adult-like decisions? To address these questions, we used intracranial stereoelectroencephalography (SEEG), a powerful tool that offers high temporal and spatial resolution for recording neural signals [39, 40, 41]. SEEG allows for precise recording from both cortical and subcortical regions, including deep brain areas often inaccessible with non-invasive methods [42, 43, 44]. This capability is critical for understanding the roles of deep brain areas in cognitive processes, which are often inaccessible with non-invasive techniques. Additionally, SEEG recordings provide robust evidence of the connectivity between cortical and subcortical brain regions[45, 29], which can further elucidate the specific network mechanisms involved in moral punishment. Building upon existing frameworks and our unique methodological advantage, we hypothesized that adults would demonstrate a more sophisticated integration of both inferred intention and punishment efficacy during TPP, supported by dynamic interactions among the vmPFC, amygdala, insula, and IPL. In contrast, we expected children to show reduced sensitivity to intention and efficacy, as well as weaker functional connectivity across these regions. We further predicted that both cognitive and neural markers of TPP would develop gradually with age. Furthermore, to gain deeper insights into the dynamic adaptation of neural circuits, we compared neural dynamics during the active task state with the intrinsic resting state, positing that processing complex contextual information in moral decision-making necessitates a fundamental shift in neural circuit configuration away from its default mode.

To test these hypotheses, 31 epilepsy patients (17 adults and 14 children) completed a novel TPP task (see Fig. 1a) during SEEG recordings to capture deep brain neural signals (see Fig. 1b). The results largely support our hypotheses. Specifically, we observed distinct broad-band gamma activity (BGA, 40-150 Hz) in the amygdala and vmPFC encoding inferred intentions in TPP decisions, with notable developmental differences in activity patterns between the adult and child patients. Moreover, vmPFC, insula, and IPL activity were critically involved in the processing and integration of punishment efficacy, also showing significant developmental variations. Moreover, connectivity analyses showed age-related variations in inter- and intra-regional communication. Specifically, the interactions among the amygdala, insula, vmPFC, and IPL differed between adults and children, and we observed evidence of intra-regional maturation within the vmPFC and IPL. Importantly, our results indicate age-dependent deviations from the intrinsic resting-state baseline in specific neural couplings (e.g., amygdala-insula, vmPFC-IPL, and vmPFC local circuits), which suggests a developmental shift in how these networks dynamically adapt to task demands. Collectively, our results revealed age-specific neural representations and connectivity patterns: while adults exhibited differentiated neural dynamics based on both intention and efficacy, children showed selective sensitivity to those cues with immature integration of outcome-related information. These findings provide novel insights into the developmental refinement of moral decision-making, which highlights the neural mechanisms that support the emergence of mature third-party punishment.

**Fig. 1.**
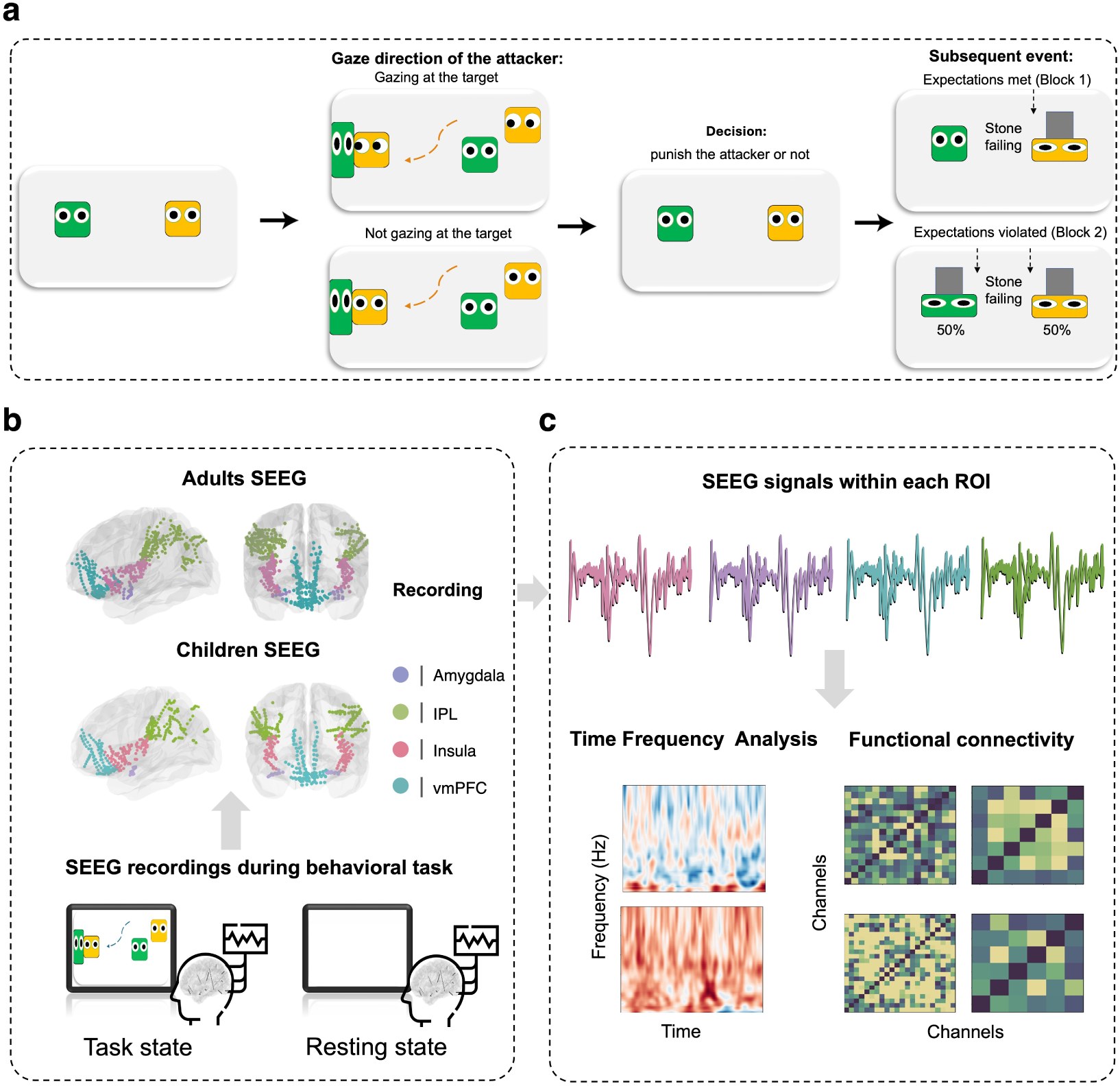
Experimental paradigm and analysis pipeline. **(a)** The behavioral paradigm was modified by Kanakogi et al.’s study[6]. Patients were first presented with a video clip of two characters interacting, where one of the two characters was attacking the other. After that, patients were asked to decide whether to punish the aggressor by releasing a stone to attack and crush it. The gaze direction of the attacker was manipulated (either following or not following the victim) to probe perceived attention. Additionally, the consequences of the punitive decision were varied in the second block, involving either crushing the attacker or the victim, enabling an exploration of the impact of expectation violation. **(b)** The general pipeline of the SEEG study. Both adult and child patients with implanted electrodes performed the task in a hospital room. Prior to the formal experiment, clear instructions and practice were provided. During the experiment, participants sat 60 cm from a computer monitor. **(c)** Raw SEEG signals were preprocessed and analyzed for different groups and experimental conditions. Time-frequency and functional connectivity analyses were subsequently performed for each group and condition.

## 2 Results

To identify the neural developmental difference in TPP, we conducted the main study on 31 epileptic patients (17 adults and 14 children) who performed the TPP task (Fig. 1a). Three adult patients were only included in the behavioral analysis because they had no available electrodes implanted in the regions of interest. A total of 28 epileptic patients (14 adults and 14 children) were involved in the following SEEG analysis (Table 1).

**Table 1.** The patients information, electrode coverage and number. vmPFC: ventromedial prefrontal cortex; IPL: inferior parietal lobule.

| SBJ | Age (years) | Gender | Group | Number of Electrodes |  |  |  |
| --- | --- | --- | --- | --- | --- | --- | --- |
|  |  |  |  | vmPFC | Amygdala | Insula | IPL |
| A01 | 31 | M | Adult | / | 7 | 2 | 8 |
| A02 | 29 | M | Adult | / | 1 | 8 | / |
| A03 | 31 | F | Adult | 3 | 6 | 9 | 17 |
| A04 | 25 | F | Adult | 15 | 5 | 7 | / |
| A05 | 31 | M | Adult | 18 | 6 | 11 | 17 |
| A06 | 20 | F | Adult | 37 | / | 10 | 8 |
| A07 | 28 | M | Adult | 20 | 3 | 12 | 8 |
| A08 | 25 | M | Adult | / | / | 8 | / |
| A09 | 25 | F | Adult | 7 | 3 | 28 | / |
| A10 | 24 | M | Adult | 5 | 2 | 6 | 34 |
| A11 | 21 | M | Adult | 17 | / | 6 | 22 |
| A12 | 20 | M | Adult | 9 | / | 2 | 21 |
| A13 | 21 | M | Adult | / | / | 10 | 39 |
| A14 | 22 | M | Adult | 6 | 4 | 3 | 6 |
| Total | / | / | / | 137 | 37 | 122 | 180 |
| C01 | 7 | F | Child | / | 3 | 11 | 16 |
| C02 | 15 | M | Child | 22 | 2 | 10 | / |
| C03 | 16 | M | Child | / | 5 | 2 | / |
| C04 | 11 | F | Child | 6 | 3 | 7 | 12 |
| C05 | 10 | M | Child | 4 | 3 | 17 | 42 |
| C06 | 16 | F | Child | / | 5 | 11 | / |
| C07 | 14 | M | Child | / | 4 | / | / |
| C08 | 7 | M | Child | / | 4 | 7 | 17 |
| C09 | 11 | M | Child | 16 | 5 | 9 | 21 |
| C10 | 12 | F | Child | 7 | 4 | 11 | / |
| C11 | 6 | F | Child | 5 | 8 | 10 | 15 |
| C12 | 14 | M | Child | 5 | 4 | 10 | 14 |
| C13 | 12 | M | Child | 50 | 5 | 2 | / |
| C14 | 10 | M | Child | / | 4 | 4 | 6 |
| Total | / | / | / | 115 | 59 | 111 | 143 |

The main task was a modified paradigm from Kanakogi et al.’s study [6]. The current task included 48 trials in total, which were divided into two blocks (24 trials each). The task started with a video clip of two characters interacting with each other, where one of the two characters was attacking the other by crushing it to the wall. Then participants were asked to decide whether to punish the aggressor by releasing a stone to attack and crush it. The gaze direction of the attacker was manipulated in the video (either following or not following the victim) to indicate harmful intention. Additionally, the result of the punitive decision was varied in the second block. When the patients chose to punish the attacker, there was a 50% chance of crushing the attacker and 50% of crushing the victim. This manipulation was to investigate the impact of punishment efficacy, which was only involved in the second block.

### 2.1 Manipulation check of the paradigm

The current experimental paradigm was adapted from Kanakogi et al.’s study [6] with additional manipulations of inferred attacking intention and the punishment efficacy. We conducted extra ratings with two independent samples of healthy adults (Exp. 1: n = 40, age = 21.08 ± 1.58 years old; Exp. 2: n = 31, age = 21.5 ± 1.70 years old) to validate the task. Meanwhile, as the youngest children involved in the SEEG sample was at the age of 5, We also did additional Experiment 3 on a healthy children sample to ensure they did fully understand and perform the task (Exp. 3: n = 11, age = 5.52 ± 0.68 years old).

In the current task, participants were expected to perceive a higher level of harmful intention of the attacker during the interaction in the intentional condition. In this case, validation ratings were added in Exp. 1 and Exp. 2 to assess participants’ feelings induced by the task. The general experimental setups for Exp. 1 and Exp. 2 were identical, where the only difference was the presenting stage of the ratings. In Exp. 1, participants were asked to rate the perceived harmful intention and their angry level on the attack after completing the whole task (Fig. S1 a, details in the supplement). In Exp. 2, participants were asked to do both ratings right after the interaction video clip in each trial, before decision-making. There was also an extra rating on the satisfaction level of the punishment results in block 2 of Exp.2 (Fig. S1 b, details in the supplement). The settings of Exp. 3 was completely identical to Exp. 1.

We first analyzed the validation ratings from Exp. 1 and Exp. 2. The results showed that participants perceived a higher level of harmful intention in the intentional condition than in the less intentional condition (Exp. 1: *t* = 4.28*, p <* 0.001***, Exp. 2: *t* = 7.16*, p <* 0.001***, Fig. S2 a). Meanwhile, the participants also reported a higher level of anger after watching the video where the attacker was considered intentional to harm the target compared to the less intentional condition (Exp. 1: *t* = 4.84*, p <* 0.001***, Exp. 2: *t* = 6.94*, p <* 0.001***, Fig. S2 b). We then compared the satisfaction ratings of the results of Exp. 2. The paired sample t-test showed that participants were more satisfied with the result when the punishment results met their expectations compared to the condition where the results violated their expectations (*t* = 6.78*, p <* 0.001***, Fig. S2 d). Overall, these results validated the TPP paradigm in the current study.

### 2.2 Behavioral performance of adults and children

We then analyzed the effect of intention on the TPP decision in Exp. 1-3. Participants were more likely to punish the attacker in the intentional condition compared to the less intentional condition in both Exp.1 (*t* = 3.76*, p <* 0.001***, Fig. S2 c left) and Exp. 2 (*t* = 6.25*, p <* 0.001***, Fig. S2 c middle). Healthy children also showed a higher probability of punishing in the attentional condition compared to the less intentional condition (*t* = 2.85*, p <* 0.05*, Fig. S2 c right). Next, we tested whether punishment results in the previous trial affected decisions in the next trial. We compared participants’ punishment decisions and their reaction time (RTs) in making decisions with different punishment results in the last trial (expectation-met, expectation-violated, and non-punished trials). Repeated ANOVA results showed that participants’ punishment rates were significantly different across the three conditions in Exp. 2 (*F* = 10.0*, p <* 0.001***). Post-hoc analysis showed that participants were more likely to punish the attacker when their expectations were violated in the last trial compared to the trials where their expectations were met (*t* = 2.61*, p <* 0.05*), and the trials where participants did not choose to punish in the last trial (*t* = 4.04*, p <* 0.001***).

Linear regression models were then used to assess the influence of intention and punishment efficacy on decision in both adults and children. To eliminate the possibility of learning effect on decision-making, we first fitted a behavioral model (M0) that only included trial and block numbers. The results showed that, except for the effect of block in the healthy adult sample, neither trial nor block significantly contributed to decision-making in any experiment (Table S1, details in the supplement). To examine whether perceived intention and punishment efficacy could predict participants’ decisions, we employed a linear regression model (M1). This model included two primary predictors: attacking intention and punishment efficacy, with reaction times (RTs), trial number, and block as control variables. Attacking intention was coded as 1 if the attacker was paying attention to the target (intentional) and 0 if the attacker was not paying attention (less intentional). Punishment efficacy was coded based on the outcome of the previous trial: 1 if the punishment expectations were met (the punishment was effective), −1 if the expectations were violated (the punishment was ineffective), and 0 if no punishment was delivered. The model was applied to each healthy participant in Exp. 1, 2, and 3. The results showed that the inferred intention (Table S2, details in the supplement) can predict the decision in the healthy adults (averaged *β* = 0.144, 95%*CI* = [0.07, 0.22]*, t* = 3.70*, p <* 0.001***). Further, we also found that the punishment efficacy (Table S2) can predict the decision in the healthy adult (averaged *β* = *−*0.02, 95%*CI* = [*−*0.05*, −*0.002]*, t* = *−*2.20*, p <* 0.05*). The inferred intention can also predict the decision in the healthy children (averaged *β* = 0.423, 95%*CI* = [0.13, 0.74]*, t* = 3.23*, p <* 0.01***). However, the punishment efficacy did not significantly contribute to children’s decisions (Table S2).

We then move on to the data collected from the SEEG patients. The one-sample t-test results revealed that the probability of punishment was significantly above chance only in the children group (*t* = 3.51*, p <* 0.01**, Fig S3 a), while the adults’ punishment rate did not significantly exceed chance levels (*t* = 1.09*, p >* 0.05). Moreover, the comparison between adult and child groups reached a marginal significance (*t* = 1.84*, p* = 0.076). This may suggest that adults’ choices are more influenced by additional factors that requires more consideration. Regarding the effect of intention, punishment rates in the intentional condition showed marginal significance in the adult group compared to the less intentional condition (*t* = 1.75*, p* = 0.099, Fig. S3 c), which was in line with the pattern observed in healthy adult controls. In contrast, child patients exhibited a significant difference between conditions (*t* = 6.25*, p <* 0.001***, Fig. S3 d). We also fitted the behavioral models with the SEEG sample. The results showed that the inferred intention (Fig. 2a) can predict the decision in adults(averaged *β* = 0.126, 95%*CI* = [0.02, 0.23]*, t* = 2.46*, p <* 0.05*). Further, we also found that the punishment efficacy (Fig. 4a) can predict the decision in adults (averaged *β* = *−*0.094, 95%*CI* = [*−*0.17*, −*0.01]*, t* = *−*2.53*, p <* 0.05*). However, neither intention (averaged *β* = 0.04, 95%*CI* = [*−*0.11, 0.20]*, t* = 0.56*, p >* 0.05) nor punishment efficacy (averaged *β* = *−*0.056, 95%*CI* = [*−*0.17, 0.57]*, t* = *−*1.06*, p >* 0.05)were significant contributors to children’s decisions in the SEEG sample (Table S2, details in the supplement). To further examine whether the influence of intention and punishment efficacy on individual decision-making increases with age, Spearman correlations were conducted to assess the relationships between model estimates for these two factors and patient age. While the correlation coefficients did not reach statistical significance, likely due to the limited sample size, we observed a trend suggesting that the impact of both intention (*ρ* = 0.27*, p* = 0.13, Fig S4 a, details in the supplement) and punishment efficacy (*ρ* = *−*0.25*, p* = 0.17, Fig S4 b) tend to increase with age.

**Fig. 2.**
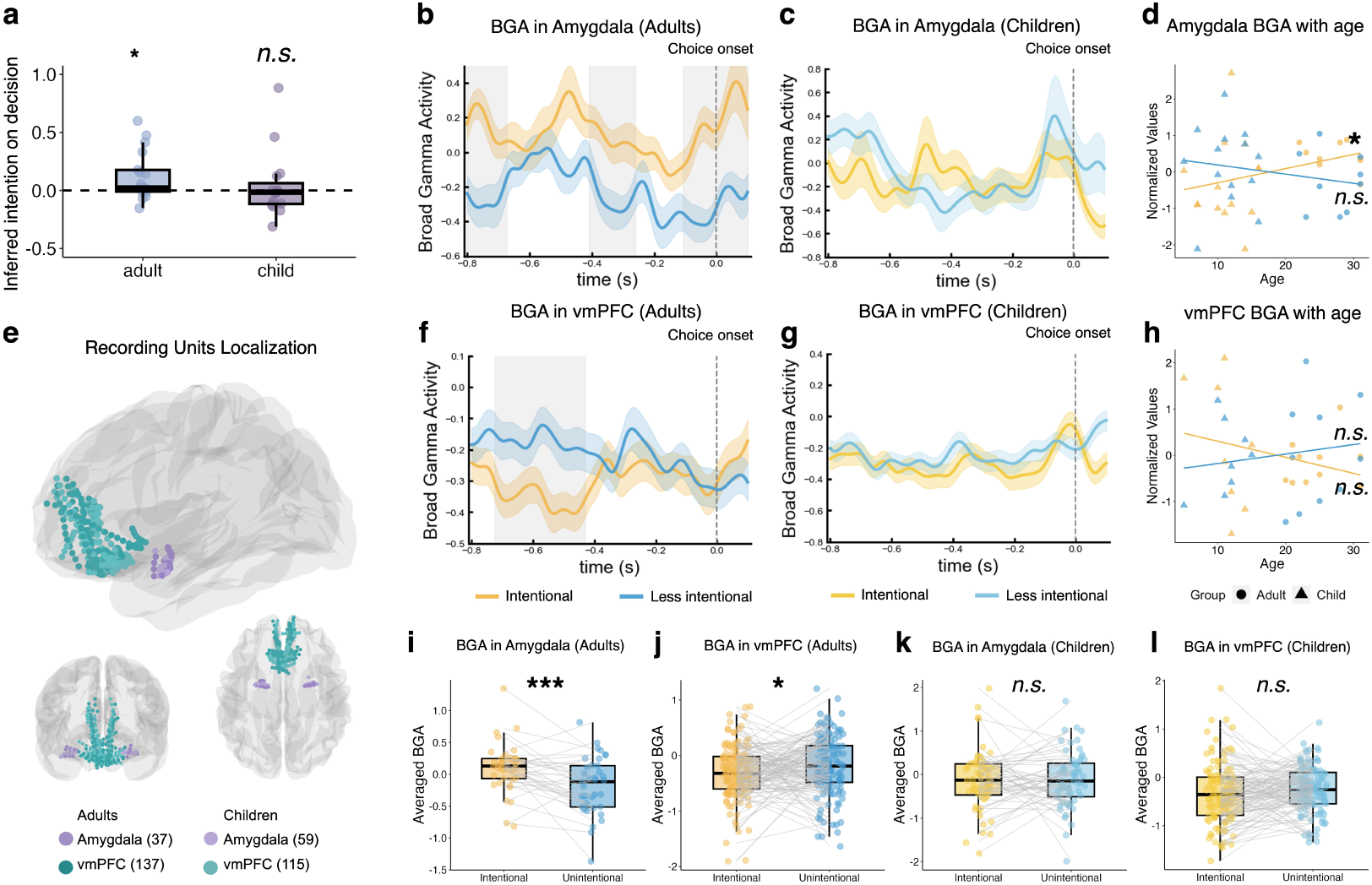
The inferred motives modulate neural activities in the amygdala, and vmPFC. **(a)** Behavioral model estimates from SEEG patients. The model included two factors: attacking intention and punishment efficacy, while controlling for reaction times (RTs) and block effects. Each dot represents the model estimate for the effect of the attacker’s intention on individual patient decisions. Results indicated that the attacker’s intention significantly influenced decision-making only in the adult group. Statistical Significant one sample t test (*p <* 0.05) result against 0 were denoted by *. **(b)** The BGA in the amygdala before choice onset in the adults group. The yellow and blue lines indicate neural activity during decision-making in the intentional and less intentional conditions, respectively. The gray shades in the background indicate the period when there was a significant difference in neural activity between the two conditions. Error ribbons represented the standard error of the mean. The zero point of the time series is the moment when the patients make the punishment decision. **(c)** The BGA in the amygdala of intentional and less intentional conditions before choice onset in children group. **(d)** Correlation between amygdala broadband gamma activity (BGA) and age. Triangles represented averaged BGA power during decision-making for child patients, while dots represented averaged BGA power for adult patients. **(e)** The anatomical location of recording sites located in the amygdala (purple), and vmPFC (turquoise). Lighter dots indicate recording sites for children. **(f)** The BGA in the vmPFC before choice onset in the adult group. **(g)** The BGA in the vmPFC before choice onset in the child group. **(h)** Correlation between vmPFC BGA and age. **(i)-(l)** The averaged BGA in the amygdala and vmPFC before choice onset in the adult and child group. * denoted significant one sample t test (*p <* 0.05) against 0, ** denoted test results *p <* 0.01, and *** denoted test results *p <* 0.001.

### 2.3 Distinct neural activation patterns underlie different inferred intentions

To investigate neural patterns during decision-making in SEEG patients facing attackers with different inferred intentions, we extracted broadband gamma activity (BGA, 40-150 Hz) from electrodes within specific regions of interest (ROIs): the amygdala (37 channels in adults, 59 channels in children), insula (122 channels in adults, 111 channels in children), and vmPFC (137 channels in adults, 115 channels in children) (see Fig. 2d and Table 1) to assess local neural activity [46, 29, 45]. Electrode locations were determined using the Brainstorm package [47] in MATLAB and verified by clinicians. We used paired sample t-tests to evaluate significant differences in BGA power across different experimental conditions. To address multiple comparisons within each time series, we used cluster-based permutation analysis and retained only the statistical test results within significant time clusters after 10,000 permutations. In adults, higher BGA was observed in three time clusters within the amygdala after the permutation test during decision-making in intentional trials compared to less intentional trials (cluster 1: −0.8 to −0.67 s from decision, *p <* 0.05*; cluster 2: −0.41 to −0.28 s from decision, *p <* 0.001**; cluster 3: −0.11 to 0.10 s from decision, *p <* 0.001**, Fig. 2b). However, a similar activation pattern was not observed in the children group (Fig. 2c).

Interestingly, distinct activation patterns were observed from the BGA within the vmPFC. Higher BGA were found before decision-making in the less intentional trials compared to the intentional trials within vmPFC in adults. After the permutation test, one significant clusters before choice onset were identified (cluster 1: −0.73 to −0.43 s from decision, *p <* 0.01**,Fig. 2f). Similar activation pattern was not observed in the children group (Fig. 2f). Although the insula is a well-documented region involved in emotion processing and affective decision-making, two significant clusters observed in the adult group did not survive permutation testing (Fig. S5; see Supplementary Information for details). Similarly, no significant differences were found in the children group.

To further investigate the neural activity before the TPP choice onset. We also calculated the averaged BGA power before decision (−0.8s - 0.1s) in each ROI. The results were consistent with the time frequency analysis. The averaged BGA power in the intentional condition was significantly higher than the less intentional condition within the amygdala in adults (*t* = 3.51*, p <* 0.001***, Fig. 2 i). Conversely, in the vmPFC, the less intentional condition showed significantly higher averaged BGA power than the intentional condition (*t* = *−*2.12*, p <* 0.05*, Fig. 2 k), but this was not observed in children (*t* = *−*0.06*, p >* 0.05*, Fig. 2l). These results suggest that the amygdala and vmPFC are involved in coding attackers’ harmful intentions during TPP decision-making. Comparing adults and children, the consideration of intention into the decision-making process may be specifically related to the functioning of the amygdala and vmPFC. To further investigate how individuals’ ability to evaluate the attacker’s intention develops with age, we calculated the averaged BGA for each patient and performed a Spearman correlation analysis with patient age. A significant positive correlation was observed between amygdala BGA and age specifically in the intentional condition (*ρ* = 0.47*, p <* 0.05, Fig. 2 d).

### 2.4 The role of the insula and vmPFC on punishment efficacy

To explore the effect of punishment efficacy, we first analyzed neural activity when patients were presented with different punishment outcomes. We focused on the same ROIs mentioned earlier: the amygdala and insula, which are related to emotion processing, and the vmPFC, involved in reasoning and value coding. We examined both adult and child patients when their expected punishment outcomes were either met or violated, which were defined as the trials with or without punishment efficacy. The results of the time-frequency analysis showed significant differences in BGA power within the insula between trials with punishment efficacy and trials without. Higher BGA in the insula was detected when they have punishment efficacy for the adults group(cluster 1: 0.34 to 0.44 s after the punishment starts, *p <* 0.05* 3b). Meanwhile, a similar effect was detected in child patients (cluster 1: 0.03 to 0.28 s after the punishment starts, *p <* 0.05*; cluster 2: 0.43 to 0.96 s after the punishment starts, *p <* 0.01**, Fig. 3c). However, distinct activation patterns were detected in the vmPFC. Specifically, the loss of punishment efficacy induced a significantly higher BGA power within the vmPFC in the adult sample compared to the expectation-met condition (cluster 1: 0.18 to 0.45 s after the start of punishment, *p <* 0.05*; cluster 2: 0.47 to 0.71 s after the start of punishment, *p <* 0.05**; cluster 3: 0.84 to 1.1 s after the start of punishment, *p <* 0.01**, Fig. 3e). Meanwhile, similar effect were also detected in the children sample (cluster: 0.40 to 0.60 s after the punishment starts, *p <* 0.05*, Fig. 3f). We also calculated the averaged BGA power after the results presentation (0s - 1.2s) in insula and vmPFC. The results were consistent with the time-frequency analysis. The averaged BGA power in the trials with punishment efficacy was significantly higher than trials without within the insula in adults (*t* = 2.11*, p <* 0.05*, Fig. 3h) and children (*t* = 2.89*, p <* 0.01**, Fig. 3i). Meanwhile, the averaged BGA power in the less intentional condition was significantly higher than the intentional condition in the vmPFC in adults (*t* = *−*3.19*, p <* 0.01*, Fig. 3f), but not in children(*t* = 0.48*, p* = 0.63, Fig. 3i). We also did not observe any significant difference in the averaged BGA after result presentation within amygdala between the two condition in neither the adult and the children sample (Fig. S4, details in the supplement).

**Fig. 3.**
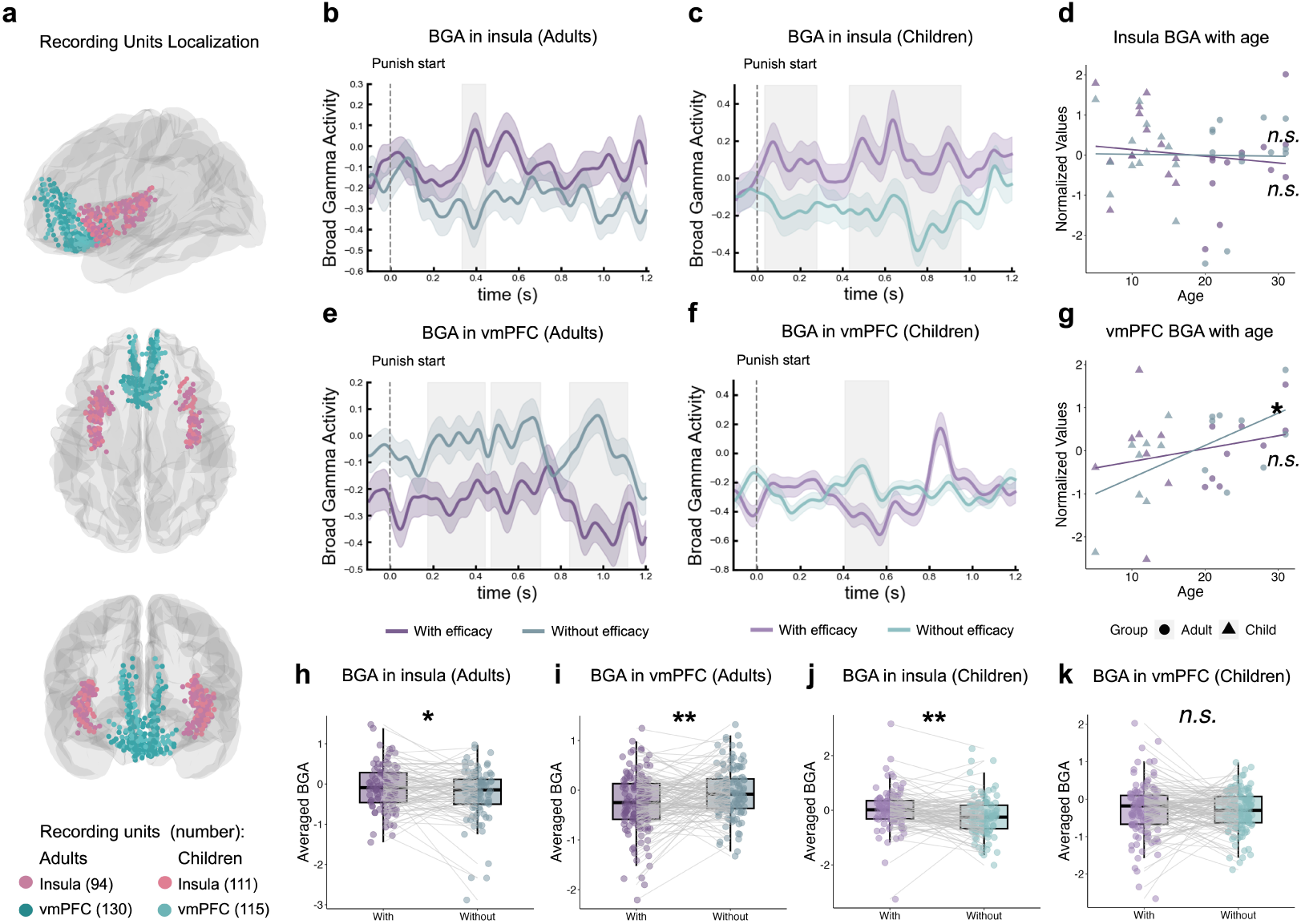
The effect of punishment efficacy on neural activities in the insula and vmPFC. **(a)** Anatomical locations of recording sites in the insula (pink) and vmPFC (turquoise). Lighter-colored dots indicate sites recorded in children. Note that the number of channels differs from the decision phase in Fig. 2 because one adult participant (A09) was excluded from this analysis due to the absence of punishment choices in the without-efficacy condition. **(b)** BGA in the insula after the presentation of the results in the adult group, represented by purple and gray (without efficacy) lines. Gray-shaded areas indicate periods of significant differences in activity between the conditions. Error bars represent the standard error of the mean. The time series is aligned to the moment of the punishment result presentation (time zero). **(c)** BGA in the insula after the presentation of the results in the children group. **(d)** correlation between insula BGA and age. Triangles represented the averaged BGA power for child patients during decision. Dots represented the averaged BGA power for adult patients, same below. **(e)** BGA activity in the vmPFC after the presentation of the results in the adult group. **(f)** BGA in the vmPFC after the presentation of the results in the children group. **(g)** correlation between insula BGA and age. Triangles represented the averaged BGA power for child patients during decision. Dots represented the averaged BGA power for adult patients, same below. **(h)-(k)** The averaged BGA in the insula and vmPFC after the presentation of the results in the adult and children group. * denoted significant one sample t test (*p <* 0.05) against 0, ** denoted test results *p <* 0.01, and *** denoted test results *p <* 0.001.

To further investigate how individuals’ ability to process whether punishment outcomes align with their expectations develops with age, we calculated the average BGA for each patient. We performed a Spearman correlation analysis with patient age. Stronger vmPFC BGA was found to be associated with older age, specifically in trials where punishment efficacy was absent (*ρ* = 0.53*, p <* 0.05, Fig. 2 g). This finding suggests that the vmPFC may play an increasingly important role in evaluating and reasoning about unexpected or suboptimal outcomes as individuals mature.

We also investigated the role of the amygdala in this process. In the adult sample, only a small cluster difference was detected between conditions (Fig. S5 a). The child sample showed increased BGA during trials with punishment efficacy, resembling the pattern observed in the insula (Fig. S5 b). However, the average BGA between conditions did not differ significantly in either group (Fig. S5 c-d), and no significant correlation was found between amygdala BGA and participants’ age (Fig. S5 e).

### 2.5 The role of IPL in the integration of punish-result expectation

To further examine how punishment efficacy influences decision-making, we analyzed neural activity based on the punishment outcome in the previous trial. The results suggest that the IPL contributes to integrating past outcomes into current decisions. Time–frequency analyses indicated that the IPL encoded the punishment efficacy from the last trial. Specifically, the with-efficacy condition of the last trial induced a significantly higher BGA power within IPL in the adult sample compared to the without-efficacy condition (cluster 1: −080 to −0.29 s from decision, *p <* 0.01**, Fig. 4b). However, There was no such effect in the children sample (Fig. 4c). The averaged BGA power before decision (0.8 - 0.1 s) in IPL also confirmed these results. The averaged BGA was significantly higher in the with-efficacy trials compared to the without-efficacy trials in adults (*t* = *−*3.47*, p <* 0.001***, Fig. 4g), but not in children (*t* = 0.68*, p >* 0.05, Fig. 4h).

**Fig. 4.**
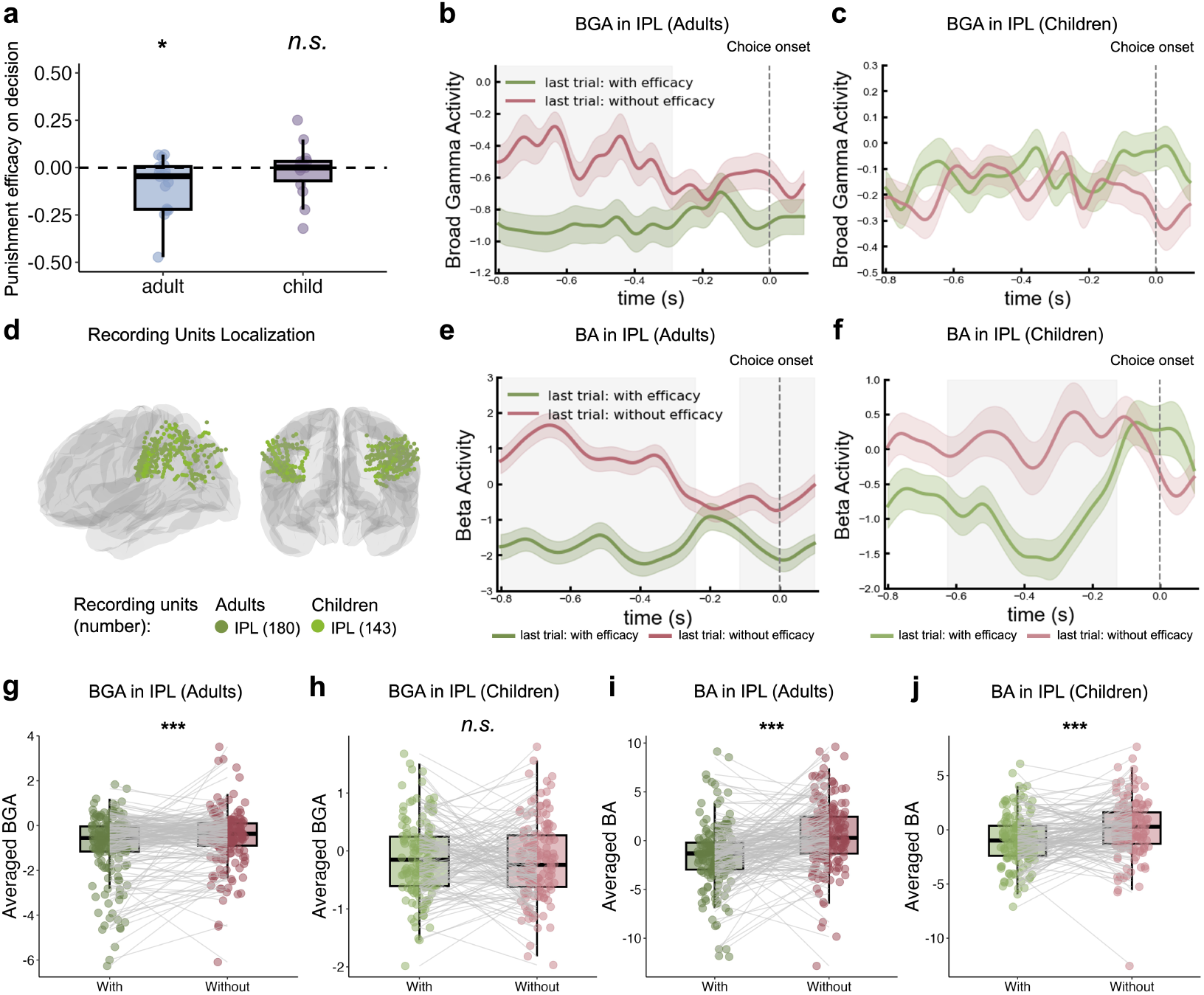
The punishment efficacy of the attacker can affect the Beta activity and BGA in the IPL. **(a)** The behavioral model estimates of the SEEG patients. The model contained two factors (attacking intention and punishment efficacy) with the reaction times (RTs) and block controlled. Significant one sample t test (*p <* 0.05) result against 0 were denoted by *. Each dot represented one patient’s model estimate of the punishment efficacy on their decision. The results suggested that the punishment efficacy was a significant contributor in the adults’ model, but not significant in the children’s model. **(b)** The anatomical location of recording sites located in IPL (green). Lighter-colored dots indicate recording sites for children. **(c)** The BA before choice onset in the adult sample. The pink and green lines indicate neural activity during decision-making when the punishment result was consistent or inconsistent with the expectation in the last trial, respectively. The gray shades in the background indicate the time window when there was a significant difference in HGA between the two conditions. Error bars represent the standard error of the mean. The zero point of the time series is the moment when the patients make a decision to punish. **(d)** The BA before choice onset in the child group. **(e)** The BGA before choice onset in the adult group. **(f)** The BGA before choice onset in the child group. **(g)-(j)** The averaged BGA and BA in the IPL during decision in the adult and children group. * denoted significant one sample t test (*p <* 0.05) against 0, ** denoted test results *p <* 0.01, and *** denoted test results *p <* 0.001.

To provide a broader view of IPL involvement, we also analysis the beta activity (BA, 12-30 Hz). The results suggested that the BA power in IPL also coded for the effect of punishment efficacy. The expectation-violated condition of the last trial induced a significantly higher BA power within the IPL in the adult sample compared to the expectation-met condition (cluster 1: −0.80 to −0.26 s from decision, *p <* 0.001***, cluster 2: −0.13 to 0.1 s from decision, *p <* 0.05*, Fig. 4 e). A similar activation pattern was also found in the children’s sample (cluster: −0.63 to −0.13 s before the choice onset, *p <* 0.001***, Fig. 4f). The averaged BGA power before decision (0.8 s - 0.1 s) in IPL also confirmed these results. Averaged BA in the 0.8–0.1 s window before decision confirmed these effects in both groups. BA was significantly higher following with-efficacy outcomes in adults (*t* = *−*6.11, *p <* 0.001***, Fig. 4i) and in children (*t* = *−*3.91, *p <* 0.001***, Fig. 4j).

### 2.6 Developmental Reorganization of Functional Connectivity during TPP Decisions

Building on the observed localized BGA changes, we further investigated the inter-regional communication among these key ROIs. We focused on phase-locking value (PLV) in the theta band (4-8 Hz), as theta rhythms are widely implicated in facilitating long-range communication and cognitive coordination across brain regions [48, 49, 45]. PLV captures precise phase synchronization, crucial for understanding task-dependent interactions between spatially distinct brain regions [50, 51, 52]. PLV between two ROIs was computed for all inter-ROI electrode pairs for each condition (Table. S3), with paired t-tests used to detect significant differences between conditions. In the current analysis, we first calculated the PLV between the insula and the amygdala to see the possible communications between the emotion processing regions before decision (Fig. 5a). Significant difference between the PLV of the intentional condition and the less intentional condition was detected in the adult patients (*t* = 2.50, *p <* 0.05*; Fig. 5b), but not in child patients (*t* = −0.81, *p >* 0.05; Fig. 5b). To further explore developmental trends, we performed a Spearman correlation between amygdala-insula PLV and chronological age. A significant positive correlation was observed for the intentional condition (*ρ* = 0.42*, p <* 0.05; Fig. 5c), which indicated an age-related increase in functional coupling during the processing of intentional harm.

**Fig. 5.**
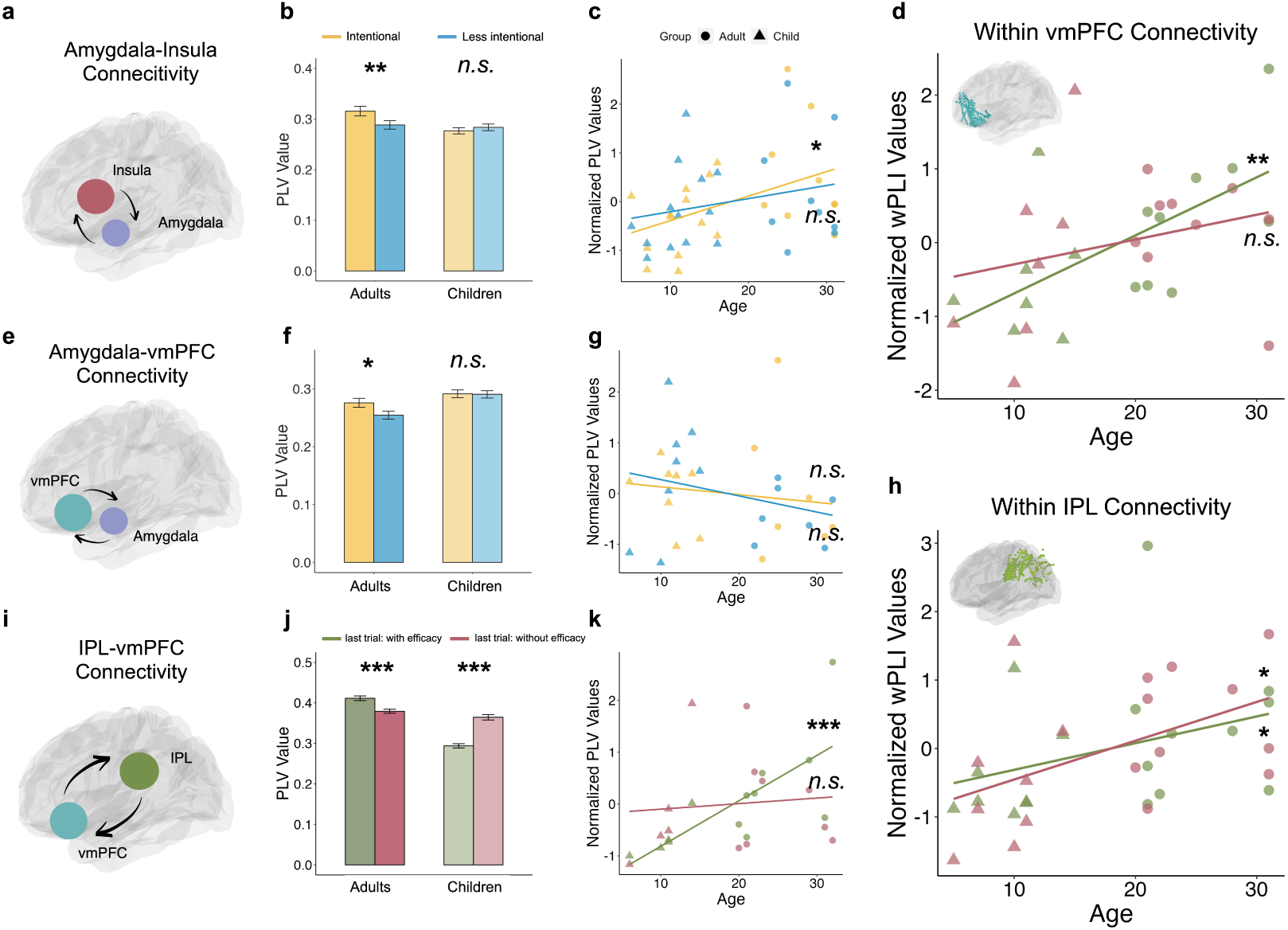
The functional connectivity during the TPP decision in adults and children. **(a)** The schematic graph of the functional connectivity between amygdala and insula **(b)** The PLV value between amygdala and insula during decision-making in different age groups’ intentional and less intentional conditions. * denoted significant one sample t test (*p <* 0.05) against 0, ** denoted test results *p <* 0.01, and *** denoted test results *p <* 0.001, same below. **(c)** The Spearman correlation between the amygdala-insula PLV and patients’ age for the intentional and less intentional conditions of the attacker’s intention. **(d)** The Spearman correlation within-vmPFC wPLI values and patients’ age in the with-efficacy and without-efficacy conditions and of different age groups. **(e)** The schematic graph of the functional connectivity between amygdala and vmPFC **(f)** The PLV value between amygdala and insula during decision-making in different age groups’ intentional and less intentional conditions. **(g)** The Spearman correlation between the amygdala-vmPFC PLV and patients’ age for the intentional and less intentional conditions of the attacker’s intention. **h)** The Spearman correlation within-IPL wPLI values and patients’ age in the with-efficacy and without-efficacy conditions and of different age groups. **(i)** The schematic graph of the functional connectivity between IPL and vmPFC. **(j)** The PLV value between IPL and vmPFC during decision-making in different age groups’ in the with-efficacy and without-efficacy conditions. **(k)** The Spearman correlation between the amygdala-vmPFC PLV and patients’ age for the with-efficacy and without-efficacy conditions and of different age groups.

To explore how can the emotional related regions and the value integration regions communicate with each other in different condition of inferred intention during decision, we also calculated the PLV between amygdala and vmPFC as well as the connectivity between insula and vmPFC under different conditions with inferred intentions. For amygdala-vmPFC connectivity (Fig. 5e), significant differences were observed in adult patients (*t* = 2.00*, p <* 0.05; Fig. 5f), with higher PLV in intentional trials, a pattern not present in children (*t* = 0.01*, p >* 0.05; Fig. 5f). However, unlike the amygdala-insula coupling, amygdala-vmPFC connectivity did not show a significant correlation with age in either condition (intentional: *ρ* = *−*0.27*, p >* 0.05; less intentional: *ρ* = *−*0.09*, p >* 0.05; Fig. 5g). Similarly, insula-vmPFC PLV did not show significant differences between intentional and less intentional conditions in either adults (*t* = 1.40*, p >* 0.05; Fig. S7) or children (*t* = *−*0.32*, p >* 0.05), nor did it correlate significantly with age (intentional: *ρ* = *−*0.11*, p >* 0.05; less intentional: *ρ* = *−*0.46*, p >* 0.05; Fig. S7). Given the role of the IPL in value integration and its functional interaction with the vmPFC during decision-making, we further examined the connectivity between these regions under the with-efficacy and without-efficacy conditions (Fig. 5i). In adult patients, PLV was significantly higher in the with-efficacy condition compared to the without-efficacy condition (PLV: *t* = 4.21*, p <* 0.001***, Fig. 5j). In contrast, child patients exhibited a markedly different pattern. Although the PLV values in children also showed a salient difference between conditions (*t* = *−*8.44*, p <* 0.001***, Fig. 5j), the effect was in another direction, where the connectivity was higher in trials without punishment efficacy. To have a deeper exploration on the differing communication patterns between groups, we calculated the Phase-Slope Index (PSI) from the vmPFC to the IPL to assess the direction of information flow under different conditions [50, 51]. The results revealed a significantly stronger flow of information from the IPL to the vmPFC in the with-efficacy condition compared to the without-efficacy condition (*t* = *−*3.15*, p <* 0.01**, Fig. S8), while no significant difference was found in the child patients (*t* = 0.85*, p >* 0.05). To further assess developmental trends, we also correlated IPL-vmPFC PLV values with age. A strong positive correlation was found in the with-efficacy condition (*ρ* = 0.87*, p <* 0.001***; Fig. 5k), but no significant correlation was detected in the without-efficacy condition (*ρ* = 0.25*, p >* 0.05; Fig. 5k). This suggests that the connectivity between IPL and vmPFC undergoes developmental refinement, specifically in conditions where punishment efficacy is evident. These results together indicate that local synchronization within vmPFC and IPL increases with age, particularly in contexts where the efficacy of punishment is salient.

Beyond interregional connectivity, we also assessed local synchronization in the vmPFC and IPL using weighted Phase Lag Index (wPLI), which is robust against volume conduction and sensitive to genuine local synchronization [53, 54]. In the vmPFC, significant age-related increases in within-region wPLI were found in the with-efficacy condition (*ρ* = 0.62*, p <* 0.01**; Fig. 5d), but not in the without-efficacy condition (*ρ* = 0.35*, p >* 0.05; Fig. 5d). Similarly, within-region IPL connectivity exhibited a significant positive correlation with age in both with-efficacy (*ρ* = 0.47*, p <* 0.05*; Fig. 5h)and the without-efficacy condition (*ρ* = 0.54*, p <* 0.05*; Fig. 5h).

### 2.7 TPP decisions reconfigures the neural dynamics away from its intrinsic state

To investigate whether TPP decision-making reconfigures these neural couplings relative to their intrinsic baseline state, we employed an Inter-subject Representational Similarity Analysis (IS-RSA) [55, 56]. This powerful second-order approach assesses how the stable pattern of neural similarity across individuals is altered across different states, moving beyond simple comparisons of activity levels. We derived functional connectivity matrices in the theta band (4–8 Hz) for resting-state and task conditions using the weighted Phase Lag Index (wPLI), chosen for its robustness to volume conduction and spurious correlations [54, 53]. We constructed subject-by-subject Representational Similarity Matrices (RSMs) for intentional, less-intentional, and resting-state conditions, where each cell represented the Pearson correlation between a pair of subjects’ connectivity patterns (Fig. 6a, Fig. S9). Initially, we examined the representational similarity of amygdala-insula coupling across the entire cohort. A bootstrapping analysis showed no significant difference in representational similarity (Δ*ρ* = *ρ_intentional,res_ −ρ_less_*_−_*_intentional,res_*) between the resting state and either the less-intentional state or the intentional state (Δ*ρ*, *p >* 0.05, 95%*CI* = [*−*0.245, 0.126]; Fig. 6b). This suggests that, at the whole-sample level, the amygdala-insula coupling did not show an overall task-evoked reconfiguration from its intrinsic state.

**Fig. 6.**
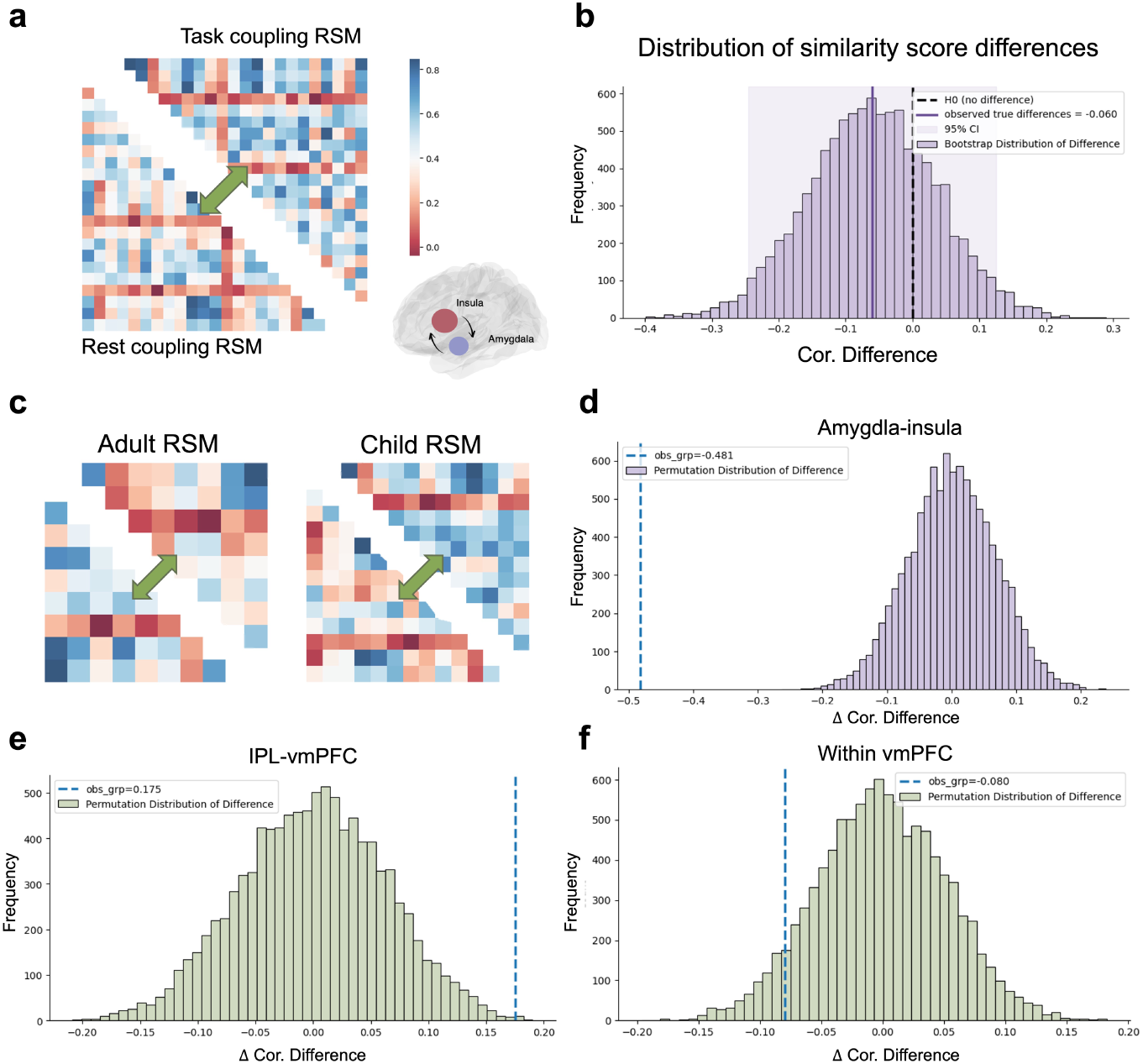
The neural reconfiguration during TPP compared to the resting state. **(a)** Intersubject representational similarity analysis (IS-RSA) pipeline. This analysis correlates the inter-subject variability in task-evoked connectivity with the inter-subject variability in resting-state connectivity. **(b)** Bootstrap distribution of similarity score differences. Each panel displays the results of a non-parametric permutation test (10,000 permutations) designed to assess significant differences. The plot shows the null distribution (histogram) and the observed group difference (dot line) for task-related network modulation. **(c)** Schematic illustration of separating the subject-by-subject Representational Similarity Matrices (RSMs) for separated adult and child groups. **(d)-(f)** Developmental interactions confirmed by non-parametric permutation testing. The test compares the degree of network reconfiguration (i.e., the difference in similarity to resting-state for task-related network modulation) between adults and children. Significant differences were found in the amygdala-insula circuit for intentional vs. less-intentional conditions between adults and children (d), and in the IPL–vmPFC circuit (e) but not in the local vmPFC processing for with vs. without efficacy conditions(f).

Given the significant developmental differences observed in previous analyses of localized BGA and functional connectivity, we next investigated whether the magnitude of this network reconfiguration differed between age groups. We separated the task-resting RSMs into adult and children groups (Fig. 6c). The permutation test was then employed to determine the statistical significance of the group interaction(Δ*ρ_adults_* = *−*0.442, 95%*CI* = [*−*0.847*, −*0.013]; Δ*ρ_children_* = 0.039, 95%*CI* = [*−*0.280, 0.357]), specifically by computing the observed difference in representational similarity scores (Δ*ρ_adults_ −* Δ*ρ_children_*). The permutation test revealed a significant group interaction for the amygdala-insula coupling (Fig. 6d). Specifically, the task-evoked reconfiguration of this functional connection by inferred intention was significantly stronger in adults than in children (observed difference of Δ*ρ* = *−*0.481*, p <* 0.001***). This indicates that in adults, processing intentional harm drives the amygdala-insula circuit into a state that diverges more substantially from its baseline intrinsic state.

Building upon the permutation test framework, we investigated developmental interactions in how punishment efficacy modulated two distinct neural aspects: inter-regional connectivity (IPL–vmPFC) and local, intra-regional processing (vmPFC). For each, we quantified this modulation as Δ*ρ* = *ρ_with_*_−_*_efficacy,res_ − ρ_without_*_−_*_efficacy,res_*. The functional connectivity between the IPL and vmPFC was significantly less reconfigured by punishment efficacy in adults (Δ*ρ_adults_* = *−*0.04, 95%*CI* = [*−*0.931, 0.824]) compared to children (Δ*ρ_children_* = *−*0.218, 95%*CI* : [*−*0.956, 0.719], observed difference of Δ*ρ* = 0.175*, p <* 0.01**, Fig. 6e). In contrast, the developmental difference in the reconfiguration of local vmPFC processing by punishment efficacy did not reach statistical significance (observed difference of Δ*ρ* = *−*0.080*, p* = 0.12, Fig. 6f). This suggests that the maturation of the IPL-vmPFC cognitive circuit involves a more stable processing state, where information integration for punishment decisions leads to representations more similar to its baseline.

## 3 Discussion

Third-party punishment (TPP) reflects a sophisticated form of moral reasoning that relies on the integration of both cognitive and emotional processes. It recruits a distributed network of brain regions whose functional organization continues to develop across childhood and adolescence. Using high-resolution SEEG recordings in both children and adults, the present study offers fine-grained spatiotemporal evidence of neural dynamics within key emotional and cognitive brain regions during TPP decisions. Furthermore, we reveal developmental changes in both intra- and inter-regional connectivity within these regions. Crucially, our findings demonstrate age-related differences in the reactive, intuitive judgments of childhood and the context-sensitive deliberations of adulthood, which is supported by the functional refinement of key subcortical and prefrontal regions.

### 3.1 Developmental differences in cognitive and emotional processing

Our current results reveal distinct behavioral and neural patterns between adult and child patients. Firstly, compared to adult patients, child patients have a higher probability of committing the punishment. These results suggest that, while children make more intuitive and direct punishment decisions, there are more contextual factors to be considered during decision-making in the adult group. Our behavioral model results are in line with this hypothesis, where inferred intention and punishment efficacy only significantly contributed to TPP decision-making in the adult patients but not in child patients. However, according to previous developmental studies, children are aware of the intent behind transgressions at the age of five [57], and the curiosity of the intent dose grow with age[58]. The results from our Exp. 3 were in line with the previous finding (Fig. S2 c), where the intention is a significant predictor of the TPP decision with the children at the age of five. However, intention did not significantly predict TPP decisions in child patients, which may relate to epilepsy-related cognitive developmental delay [59, 60]. Although we only include children with normal cognitive function, developmental delays associated with epilepsy may still impact decision-making [61]. Additionally, prolonged hospitalization and limited social interaction may also hinder moral and social learning, as emotional and social experiences play a crucial role in moral development [20, 62].

At the neural level, we find developmental differences during both the decision and outcome evaluation stages. Moral evaluation and punish behavior develop from a very young age, with a more instinctive manner. A recent study shows that TPP behavior can even be detected in pre-verbal infants [6]. It is also shown that infants’ brain responses to static pictures of a helper and hinderer indicate differential processing of social information as early as 6 months of age [7]. Despite moral principles and intuitive moral judgments taking shape early in childhood[6, 7, 4], children may still lack a deep understanding and appreciation of the moral principles. This deeper comprehension requires encoding multiple contextual and situational factors, which are crucial for more comprehensive moral decisions [8, 9, 63]. This suggests that TPP decisions in children are likely to be more reactive and driven by automatic emotional responses, whereas the capacity for deliberate and integrative moral reasoning develops progressively with age and neural maturation (Fig. 7).

**Fig. 7.**
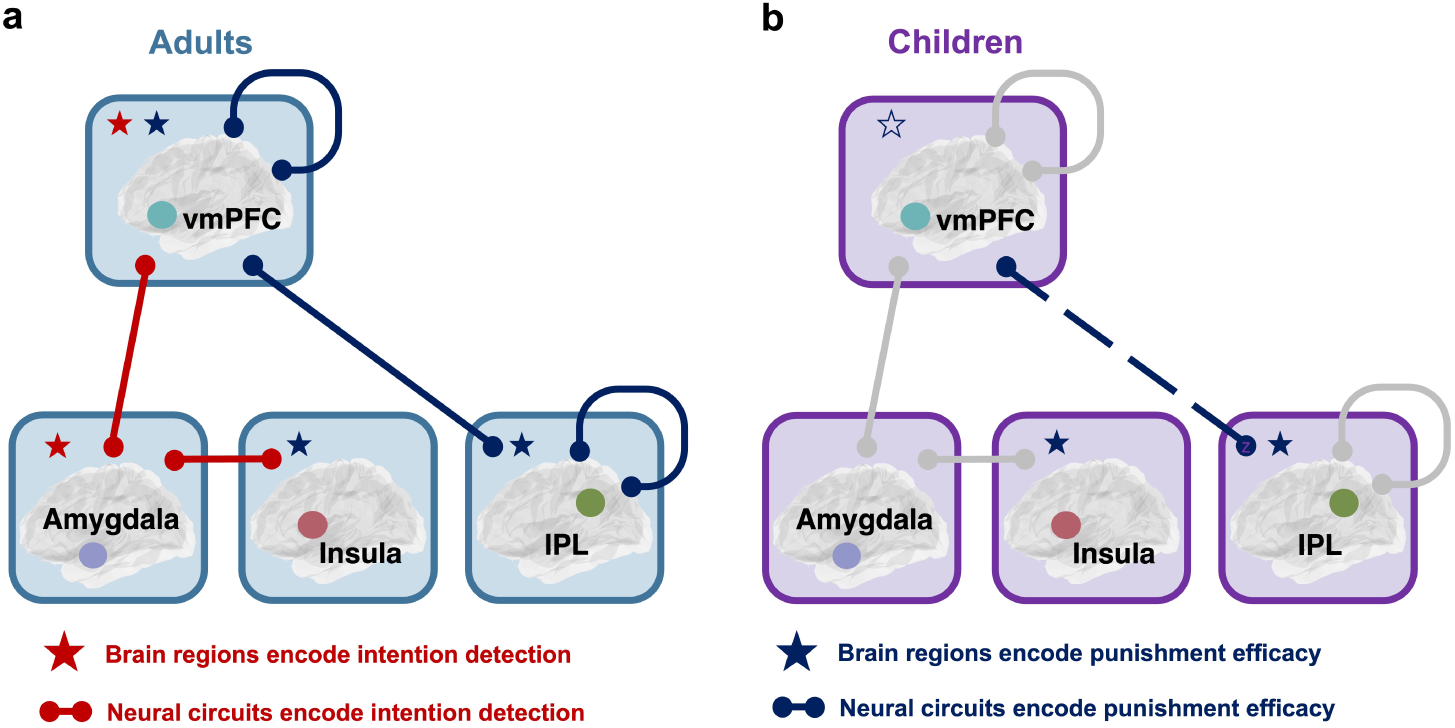
Graphical Summary. **(a)** In adult patients, mature TPP decision-making integrates both intention and punishment efficacy through the coordinated activity of the vmPFC, amygdala, insula, and IPL. Specifically, the vmPFC-amygdala-insula circuit plays a key role in detecting the perpetrator’s intention, while the vmPFC-IPL coupling is involved in evaluating the consequences of punishment. Red stars indicate regions with significant differences between intentional conditions, and red connecting nodes denote significant connectivity. Navy stars and connecting nodes indicate regions and connectivity encoding punishment efficacy. **(b)** In child patients, the neural mechanisms underlying TPP are still developing. Condition-specific activation and functional connectivity in the intention-detection circuit (vmPFC-amygdala-insula) are not yet observed. Although children exhibit similar BGA activation patterns in response to punishment efficacy as adults, the vmPFC-IPL coupling and intra-regional connectivity remain less developed, potentially limiting their ability to fully integrate punishment outcomes into decision-making. Dashed lines indicate different connectivity patterns compared to adults, while grey connecting nodes refer to non-significant connectivity between conditions in child patients.

### 3.2 The recruitment of amygdala and vmPFC in intention inference

Our results highlight that the inferred intention of the attacker can affect the TPP decision-making in adults. This aligns with the previous findings that people are highly sensitive to harmful intentions in everyday life [12]. Prior studies have found that individuals report higher levels of pain when they believe the harm was caused intentionally rather than accidentally [14], and that intentional harms consistently elicit stronger blame and harsher punishment than unintentional ones [13, 15, 16]. Our findings add a novel neural perspective to this behavioral literature by showing how the brain’s mechanisms underlying punishment decision-making vary as a function of inferred intentions and developmental stage. Specifically, we observe distinct patterns of amygdala and vmPFC activity across different conditions, highlighting their complementary roles in processing intentionality and guiding moral decision-making.

The amygdala shows significantly higher BGA during intentional harm conditions compared to less intentional conditions in adults, particularly in the time windows leading up to the decision. This finding is consistent with previous research suggesting that the amygdala plays a crucial role in encoding emotional salience and social threat perception [25, 64, 65]. Notably, this effect is absent in children, which suggests that the sensitivity of the amygdala to intentionality may develop over age, which can also be supported by the positive correlation between the amygdala BGA and patients’ age. The increased amygdala engagement in intentional harm trials likely reflects heightened emotional arousal and threat evaluation, which is further supported by our connectivity analyses. Notably, our IS-RSA results showed that perceiving harmful intent drives the amygdala–insula coupling into a functional configuration that is distinct from the brain’s intrinsic resting-state mode. This provides mechanistic evidence that intentionality triggers a reconfiguration of the salience network, mobilizing affective processing resources for moral evaluation [66, 67, 21].

In contrast, the vmPFC exhibits an opposite pattern: adults showed significantly higher BGA in response to less intentional harm. The vmPFC is well-documented for its role in integrating social and emotional information to guide context-sensitive decision-making [68, 31]. This pattern suggests that in situations where harm is evaluated as less intentional, the vmPFC may be engaged in downregulating punitive impulses, which may facilitate a controlled response rather than an emotionally driven reaction. This effect was also absent in the child patients, which further indicates developmental differences in regulatory processing within the vmPFC. Our results highlight the importance of considering both condition-specific and developmental differences in the neural mechanisms underlying social decision-making. The amygdala’s heightened sensitivity to intentionality and the vmPFC’s role in regulatory control appear to emerge with age, supporting a more sophisticated evaluation of moral and social contexts.

### 3.3 The insula and vmPFC track the punishment efficacy

Our findings also highlight significant neural differences in the processing of punishment efficacy across both conditions and developmental stages. Specifically, we observed distinct activation patterns in the insula and vmPFC when punishment outcomes aligned with or deviated from expectations. The insula showed increased broadband gamma activity (BGA) when punishment efficacy was met, suggesting its role in reinforcing the expected consequences of actions. This result aligns with prior studies emphasizing the insula’s role in affective processing and salience detection [25, 28, 69]. It has been well acknowledged that the amygdala and insula are both key regions involved in emotional processing [70, 26, 71, 25, 69]. The lack of significant amygdala activation differences between conditions in adults suggests that, while the amygdala is crucial for emotional responses, the processing of punishment efficacy may rely more on the integration of affective and cognitive signals within the insula [72]. Interestingly, children exhibit the same neural response patterns as adults when receiving punishment results, indicating similar emotional processing capabilities regarding punishment outcome processing.

Moreover, the increased BGA power in the vmPFC in response to expectation-violation was also detected, which may indicate that enhanced reasoning and emotional regulation are required during this process [73, 31]. The vmPFC has long been implicated in numerous roles across social, cognitive, and affective processes and is often disrupted in various mental illnesses [31]. The vmPFC is essential for the generation and regulation of negative emotions [31].

The differences observed between adults and children provide insight into the neurodevelopmental changes in punishment processing. Both age groups show increased insula activation when punishment expectations were met, which contradicts our hypothesis that these regions might not be mature enough to process unexpected results. This discrepancy may reflect developmental differences in how the insula and vmPFC processes emotional as well as cognitive information at different stages. While the vmPFC appears to engage similarly in both adults and children during the results-revealing stage, we find a significant correlation with the vmPFC BGA and age, especially in the without-efficacy condition. It indicates an increased reasoning about unexpected outcomes. Prior work also suggests that the vmPFC undergoes prolonged maturation, refining its role in integrating contextual information for decision-making[74]. Together, the results from the results-revealing stage provide converging evidence that adults and children show similar emotional and cognitive responses to the punishment efficacy. Consequently, these similarity indicates that although children can understand the social norms and punishment efficacy, they may still need to acquire the ability of integrating the punishment results into their further decisions as they mature.

### 3.4 The role of IPL in information processing and integration

Beyond the emotional responses elicited during the results-revealing stage, punishment efficacy also influences decision-making in subsequent trials. Behavioral models suggest that punishment efficacy is a significant contributor to TPP decisions in adult patients, but not in child patients. Additionally, distinct and shared neural patterns are observed in integrating punishment efficacy into subsequent decision-making processes in adults and children.

Neurally, adults shows higher BGA and BA power in the IPL during decision-making when prior punishment was effective, whereas child patients have similar patterns only in BA power (Fig. 4 f, j), which indicates less mature IPL functioning. These findings are consistent with previous findings that IPL engages expectation violation and memory retrieval during decision making[35]. The IPL serves as a convergence zone for various brain networks, and is central to executing key cognitive operations across different levels of neural processing [75, 76]. In this case, although both adults and children share similar neural responses during the outcome stage, the matured functioning of IPL may be specifically required in the integration of the punishment results in adults. Our current findings further elaborate on the special roles played by the IPL during social decision-making. Additionally, the results indicate that the function of the IPL develops relatively early in the lifespan compared to other cortical regions. Furthermore, inter-regional connectivity between the IPL and other cortical areas is crucial in guiding individual choices. Our FC results provide further evidence of this view, where we find significant BGA coupling from the IPL to the vmPFC exclusively in adults when their punishment efficacy is high in the previous trial. These findings together suggest that the activation of the IPL alone is insufficient for integrating punishment efficacy into current decision-making, which requires the coordinated work of the IPL and vmPFC for this process.

### 3.5 Condition-dependent functional connectivity and the developmental differences

This study provides a comprehensive framework and developmental insights into TPP decision-making. By disentangling the frequency, time, and functional significance of neural responses in four key nodes of the emotion and decision-making brain regions, our findings highlight the developmental features underlying TPP behavior. In particular, the observed condition-dependent changes in functional connectivity emphasize the importance of inter-regional communication in processing inferred intentions and punishment efficacy. Beyond simply examining connectivity strength, our analysis further investigates how task demands reconfigure neural dynamics away from their intrinsic resting state, revealing crucial developmental differences in this capacity.

Adults show enhanced amygdala-insula synchrony between regions associated with emotional processing and social evaluation when processing intentional harm. The amygdala and insula are central to affective processing and moral assessment [77, 78], and increased coupling between these areas in adults likely reflects more efficient integration of emotional and contextual cues during moral decision-making. These findings are in line with prior research demonstrating that amygdala-insula synchronization facilitates the encoding of emotional intensity and intentionality[66]. Conversely, children do not show such differences, which implies that the underlying neural circuitry may not yet be fully differentiated with respect to task demands. The age-dependent increase in amygdala-insula coupling during intentional harm processing suggests a gradual refinement in emotional network interactions, which is consistent with previous study that the connectivity between amygdala and insula is important for emotion regulation[79, 78]. Prior studies have shown that these regions undergo substantial synaptic pruning and myelination during development, facilitating more efficient emotion regulation and moral reasoning [74, 80]. Further, we also find that the magnitude of task-evoked reconfiguration was significantly stronger in adults than in children during intentional harm processing. This suggests that mature individuals’ neural networks undergo a more pronounced, emotion related state shift when confronted with intentional moral violations, enabling a more specialized processing mode.

Similar condition-dependent effect is also observed in amygdala–vmPFC connectivity, where only adults show significant differences. The vmPFC is a key hub for integrating affective signals with cognitive control, particularly in moral and social decision-making [9, 81]. The increased amygdala-vmPFC connectivity in adults when processing intentional harm suggests a greater recruitment of top-down regulatory mechanisms, which may contribute to the more contextually sensitive punishment decisions observed in mature individuals[18]. This is also in line with the previous studies that vmPFC is critical for the regulation of the amygdala and that the amygdala–insula–vmPFC network is essential for affective and social decision-making [82, 36, 78]. In contrast, children’s relatively stable amygdala-vmPFC connectivity across conditions also suggests that their judgments may be more directly driven by intuition rather than complex moral reasoning.

In terms of value-based integration, IPL-vmPFC connectivity showed a notable difference between adults and children. In adults, stronger connectivity was observed when punishment was effective, whereas children showed the opposite pattern, with higher connectivity when punishment lacked efficacy. The IPL plays a crucial role in integrating contextual and social information to guide decision-making [83, 84], and its increased synchronization with vmPFC in adults suggests a more refined assessment of punishment efficacy. This reversal of condition-related connectivity between age groups suggests that children may rely on more rigid, outcome-insensitive heuristics, while adults engage more flexible, context-sensitive evaluation strategies [63]. The age-related increase in IPL–vmPFC connectivity suggests that the capacity to weigh the consequences of punitive actions may be limited in younger individuals due to immature long-range network integration. This further supports the notion that the ability to integrate contextual value signals into moral decisions develops progressively with cortical maturation[85, 86]. Meanwhile, local connectivity within the vmPFC and IPL increased with age, particularly in conditions involving effective punishment. Previous fMRI study also suggests that vmPFC can carry contextual information that can affect future choices[87, 88, 32, 89]. These findings suggest that local information processing within key decision-related regions becomes more efficient over development, potentially enabling the transition from intuitive judgments to deliberate, outcome-sensitive reasoning.

Interestingly, IPL–vmPFC coupling was significantly less reconfigured by punishment efficacy from the resting state in adults than in children. Both vmPFC and IPL are core components of the Default Mode Network (DMN)[21, 75], which is critical for mentalizing, internal model maintenance and memory retrieval[21, 90]. Our observation of IPL–vmPFC connectivity in adults showing less reconfiguration by punishment efficacy compared to children suggests that in the mature brain, this DMN-related circuit operates with greater stability[91]. Rather than undergoing substantial representational shifts in response to task-relevant variables, the adult DMN components may maintain a more consistent functional state. This could reflect a more efficient integration of external stimuli into a robust internal model[92]. Together, the significant interactions suggest a shift towards a more mature functional state where information is integrated within a more established and resilient neural framework pf IPL and vmPFC.

In sum, our FC results show that TPP decisions rely on coordinated activity between emotional and cognitive systems, with distinct patterns emerging across development. In adults, functional connectivity between the vmPFC, amygdala, insula, and IPL is dynamically modulated by both inferred intention and punishment efficacy, which reflects a mature, context-sensitive integration of social information. In contrast, children show more stable connectivity patterns, suggesting a greater reliance on intuitive, reactive processes (Fig.7). These findings provide compelling evidence that the neural architecture supporting third-party punishment undergoes substantial developmental refinement.

### 3.6 Limitations and future directions

While our study provides high-resolution insights into the neural correlates of TPP using intracranial SEEG, we acknowledge certain limitations. First, the inherent constraints of clinical SEEG recordings, obtained from epilepsy patients, resulted in a relatively small sample size. Notably, the small sample size within the child group prevents a finer-grained classification by age, which could offer deeper insights into age-related differences in TPP decision-making and neural development. Age is a critical factor in brain maturation, particularly in the prefrontal cortex and its connectivity with subcortical regions. Therefore, the lack of detailed age subgroups within the child patient group limits our understanding of how these developmental changes unfold over time. Future studies with larger sample sizes and more refined age stratification are welcome to validate and extend our findings.

A second limitation concerns the coverage and spatial resolution of SEEG. While SEEG is a unique access to deep brain structures with high temporal and spatial resolution, electrode placement is determined by clinical needs rather than experimental design. As a result, only a subset of patients had electrodes implanted in more than one region of interest (ROI), which reduces the number of electrode pairs available for functional connectivity analyses. This constrains our ability to draw stronger inferences about network-level interactions, as well as about the specific contributions of subregions within the amygdala or insula. Meanwhile, although SEEG provides precise spatial resolution within deep brain structures, it can only be recorded in neurosurgical patients. It raises concerns about the applicability of these findings to neurologically healthy individuals. But still our findings are consistent with the previous findings using the non-invasive methods.

Despite these limitations, our findings rise several promising possibilities for future research. For instance, the observed age-related differences in TPP behavior and neural activation patterns highlight the importance of considering developmental trajectories in studies of moral cognition. Future work could explore how other factors, such as social context, cultural background, or individual differences in empathy—interact with age to shape TPP-related decision-making. In addition, considering the influence of social factors such as group membership and peer influence could enrich our understanding of moral reasoning in real-world contexts[93].

Finally, our results have potential implications from a translational perspective, they may inform debates in developmental psychology, forensic science, and legal policy, particularly regarding the age at which individuals should be considered morally and legally responsible for their actions [94]. Understanding the neural underpinnings of moral decision-making across development could help shape age-appropriate interventions and inform judicial practices involving minors.

## 4 Conclusion

In summary, our behavioral and SEEG findings demonstrate that mature third-party punishment decisions rely on the integrated work of emotional systems (including the amygdala and insula) and cognitive networks (including the vmPFC and IPL). Refined inter- and intra-regional connectivity supports this integrated processing, enabling individuals to consider both inferred intentions and contextual factors during moral decision-making (Fig. 7). In contrast, the immature interplay between these systems in children appears to favor more direct, intuitive choices that may overlook broader contextual factors. Our results provide novel intracranial evidence for the dynamic interplay between affective and cognitive processes in moral decision-making, which further highlight the importance of developmental context in shaping the neural architecture that supports moral judgment.

## 5 Materials and Methods

### 5.1 Participants

We conducted the main study on 31 epileptic patients (17 adults and 14 children). Three adult patients were excluded in the SEEG analysis because no available electrodes implanted in the ROI. 14 adult patients (4 females; age, 25.36 ± 4.27 years, mean ± SD) and 14 child patients (5 females; age, 11.50 ± 3.30 years, mean ± SD) were included in the formal SEEG analysis. The patients were implanted with intracranial depth electrodes for a presurgical evaluation of their epilepsy onset regions. The neural surgeons determined electrode placement according to the patients’ clinical diagnosis. The study has been approved by the Ethics Committee of the University of Macau (BSERE22-APP015-ICI-M1) and Xiamen Humanity Hospital (HAXM-MEC-20230915-038-01). Informed consent was obtained from all patients (and their guardians for patients under 18 years old), and all procedures adhered to the guidelines set by the clinicians and IRB regulations.

### 5.2 Experimental paradigm

The experimental design was a modified task of the paradigm from Kanakogi et al.’s study (2022). The current paradigm included a total of 48 trials, divided into two sequential blocks of 24 trials each. In the current study, patients were first presented with a 7-second video clip of two characters interacting with each other, where one of the two characters was attacking the other. After watching the video, patients were asked to decide whether to punish the aggressor by releasing a stone to attack and crush it using the key board. Specifically, on each trial they could choose to administer punishment by pressing the key “3” or refrain from punishment by pressing the key “1”. If a punishment decision was made, an outcome presentation was displayed for approximately 1.5 seconds; if the participant chose to refrain from punishment, a blank screen was shown instead.

Within each block, the intentional and less intentional conditions were interleaved using randomization to minimize order effects. In particular, the gaze direction of the attacker was manipulated within the video clips to indicate different levels of harmful intention: when the attacker’s gaze was directed at the victim, it signified an “intentional” condition; when the gaze was not directed at the victim, it indicated a “less intentional” condition. For the manipulation of punishment efficacy, in Block 1, punishment was always effective. In contrast, Block 2 introduced a manipulation of punishment efficacy. Here, when participants chose to punish the attacker, there was a 50% chance of the punishment being effective (i.e., the stone correctly hit the attacker) and a 50% chance of it being ineffective (i.e., the stone hit the victim). This manipulation was designed to examine the impact of expectation violation on decision-making. For the validation of experimental manipulation, we conducted behavioral Exp. 1 (n = 40, age = 21.08 ± 1.58 years old) and Exp. 2 (n = 31, age = 21.5 ± 1.70 years old) to check the validity of the video materials and the subjective feelings induced by the interaction scenario of the two characters across participants. In Exp. 1, participants were required to rate the level of deliberation of the attacker and their anger levels on the attacking behavior after completing the whole task. In Exp. 2, the general experimental setting was identical to Exp. 1, but participants were required to do the deliberation and anger level ratings every time they watched the interaction in each trial. Participants were also required to rate their satisfaction level with the results every time when they made the punishment decision in block 2 to make sure the manipulation of expectation violation was validated. The task used in the SEEG recordings was identical to Exp. 1 but with no ratings.

During the SEEG recording, patients with electrode implementation performed the task in the hospital room. All participants were provided with written informed consent. The consent was also obtained from the guardians of all the child patients. Before the task, participants were presented with clear experiment instructions. During the experiment, participants were seated on a hospital bed, positioned 60 cm away from a computer monitor. Following the completion of the informed consent form, the practice session started. During this practice session, participants were presented with the shorter version of the video with less interaction between the two targets. All patients reported that they fully understood the task after the practice session.

### 5.3 Behavioral analysis

To examine the role of intention and punishment efficacy played during decision-making, linear regression models were fitted for each participant in both groups to investigate the specific roles played by each factor. To eliminate the possibility of learning effect on decision-making, we first fitted a behavioral model (M0) that only included the trial number and block number for each participant. The results show that none of them were significant contributors in either adults or children groups (Table. S1, details in the supplement).

To investigate whether perceived intention and punishment efficacy could predict decisions (M1), using linear regression, we predicted subsequent decisions with two factors (attacking intention and punishment efficacy). The attacking intention was coded as whether the attacker was paying attention (= 1) or not paying attention (= 0) to the target. The punishment efficacy was coded as whether the punishment results met (= 1) or violated (= 0) their expectation of punishing the attacker (Table. S2, details in supplement). The full model included inferred intention, and punishment efficacy, controlling variables including the reaction time, trials and block.

### 5.4 Intracranial EEG recordings, localization, and preprocessing

The intracranial recordings were collected using SEEG monitoring in patients undergoing clinical treatment for drug-resistant epilepsy [95]. Electrophysiological signals were acquired from electrodes implanted directly into the brain parenchyma, with a sampling frequency of 2000 Hz. The signals were collected from standard clinical penetrating depth electrodes (Sinovation, China) implanted into the brain parenchyma. Each electrode has of 8–16 contacts, with a diameter of 0.8 mm for the depth electrode. The length of each contact was 2 mm, and the distance between contacts was 1.5 mm. The trajectory for all planned electrodes was prepared using the manufacturer’s ROSA planning software (version 2.5.8) by loading and fusing MRI and CT scans. Pre-implantation MRI with trajectory plans was merged with post-implantation CT scans obtained with electrodes in place to assess the accuracy of the electrode positions relative to the preoperative planned trajectories. The number and placement of electrodes were determined by neurosurgeons for each patient, aiming to localize epileptogenic brain regions as part of the evaluation for epilepsy surgery. Task events in the SEEG recordings were marked by the TTL pulses generated in the computer and sent to a parallel input port designed for the SEEG signal amplifier (Neuralynx Inc., Bozeman MT).

The Brainstorm package [47] in MATLAB was used to label and localize the electrode contacts. For each patient, presurgical anatomical T1 MRI and post-operative CT images with SEEG electrodes were acquired. MRI segmentation was conducted using the CAT12 toolbox, facilitating the delineation of brain structures and tissue types. Subsequently, the patient’s MRI scans were spatially normalized to a standard MNI 152 template brain using the SPM mutual information algorithm, enabling group-level analyses across participants. The post-operative CT images, containing the SEEG electrodes, were co-registered to the MNI space using the SPM12 toolbox (https://www.fil.ion.ucl.ac.uk/spm/software/spm12/). Manual identification of the electrode locations was performed directly on the co-registered CT images to accurately localize each electrode within the brain volume. Three parcellations were used for labeling each recording site. AAL3 parcellation [96] was used for labeling electrodes in the insula and amygdala. The Human Brainnetome Atlas was used to localize IPL, STS, and ACCg [97]. Additionally, electrodes in the vmPFC were labeled using a vmPFC mask available at https://neurovault.org/images/18650/.

SEEG data preprocessing was conducted using the EEGLAB toolbox [98] in MATLAB (R2023b). Raw SEEG recordings underwent meticulous manual inspection by either a neurologist or an experienced clinical SEEG data analyst to identify channels exhibiting epileptic spiking or other forms of artifact contamination were excluded from the analysis. Channels containing such activity were excluded from subsequent analyses to ensure data integrity. The preprocessing pipeline, based on our previous work [45], involved downsampling the data from 2000 Hz to 1000 Hz, applying a Butterworth bandpass filter from 1 to 150 Hz, and re-referencing the data in a bipolar montage with neighboring channels to improve spatial specificity and reduce noise [42, 99, 51]. Additionally, a bandstop filter was applied at 50 Hz and 100 Hz, using a Butterworth filter with a bandwidth of 2 Hz, to effectively remove line noise and its harmonics from the SEEG recordings. Then the preprocessed data were segmented into individual trials, each time-locked to the onset of stimuli. The epoch duration spanned from 1.0 seconds prior to the stimulus onset to 4 seconds post-stimulus, which can facilitate the isolation and analysis of neural responses elicited by the experimental stimuli while accounting for both pre-stimulus and post-stimulus activity dynamics. For the resting-state data, the preprocessing pipeline was identical to the task data. After preprocessing, the continuous recordings were first segmented into non-overlapping epochs of 2.0 seconds. To isolate the frequency band of interest, a finite impulse response (FIR) filter was applied to band-pass the epoched data into the theta range (4–8 Hz) for future analysis.

### 5.5 Time-frequency feature extraction

Beta band neural activity (BA) and broad-band gamma neural activity (BGA) were extracted for analysis, which was selected based on previous studies indicating broadband activity spatially precise measure of local neuronal population spiking [100, 42, 101]. We extracted the BA and BGA using the MNE module [102] in Python. The frequencies were extracted using the Morlet wavelet transform [103] to decompose the epoch data into 12-30 Hz and 40-150 Hz respectively [104, 105]. For the ROI-based analysis, the time-frequency signals for all trials under the same conditions (e.g., attention vs, non-attention) were averaged for each electrode. Subsequently, baseline correction was implemented for each frequency band by dividing the power at all time points within a trial by the average baseline power, which was 120 ms before the cartoon before they started to move, and applying a logarithmic transformation (log-ratio) following the method outlined by [106]. To create a one-dimensional signal, frequencies within the BA and BGA were averaged, generating a single BA and BGA time course per electrode. Lastly, low-pass filtering using a Gaussian window was conducted (width = 0.1 s) for further analysis. After extracting the BA and BGA for within-ROI analysis, a value matrix with dimensions of [Electrodes * Time points] across all participants was obtained for each condition.

### 5.6 Within-ROI analysis

In our study, we conducted a within-ROI analysis to investigate the significance of neural activation between two conditions, as well as its difference from the baseline [107]. We used paired-sample t-tests to evaluate the significance of neural activation between the two conditions. Meanwhile, considering the possible issue of comparing multiple results and ensuring the reliability of our findings, we used a cluster-based permutation test [108]. This method is well-suited for analyzing time series data as it considers the temporal closeness by grouping time series. We set the p-value threshold for permutation at 0.05 and performed 10,000 permutations to strengthen the validity of our analysis.

### 5.7 Functional Connectivity analysis

In this study, we investigated functional connectivity (FC) between different regions of interest (ROIs) (vmpfc, IPL, insula, and amygdala) using SEEG data. To ensure that our results were not biased by the choice of metric, we used multiple connectivity measures for different purposes of the analysis: phase-locking value (PLV), weighted phase lag index (wPLI), and phase-slope index (PSI). Previous research has extensively demonstrated that the coupling of theta rhythms facilitates communication across brain regions [48, 49, 45]. Therefore, in the present study, our FC analysis focused on the low-frequency range within the theta band (4-8 Hz). All the FC related analysis was performed using MNE-Connectivity module from python[102]. For each subject, epoch data was extracted for each ROI. A bandpass filter for the theta band (4–8 Hz) was applied to each ROI’s data using a finite impulse response (FIR) filter to preserve phase information. Data from the two ROIs were then concatenated along the channel dimension to compute connectivity for every channel pair between ROI1 and ROI2, which that all possible interactions between channels from the two ROIs were considered. Then the different metrics were calculated for each pair of channels.

#### 5.7.1 Phase Locking Value

Phase-locking value (PLV) is also a widely used metric in EEG and intracranial EEG studies to quantify the synchronization of neural activity between different brain regions. PLV measures the consistency of phase differences between oscillatory signals recorded from distinct electrodes over time, which can provide an index of functional connectivity across channels. PLV values range from 0 (indicating no synchronization) to 1 (indicating perfect phase alignment). Because PLV focuses solely on the timing relationships between signals, it is robust against variations in signal power and noise, which is a feature particularly advantageous for SEEG recordings from deep brain structures with high temporal resolution. This makes PLV particularly useful for evaluating inter-regional connectivity where reliable phase relationships are critical [50, 52].

#### 5.7.2 Weighted phase lag index

The phase lag index (PLI) quantifies synchronization between neural signals while minimizing the influence of volume conduction or common sources. PLI focuses on the asymmetry of phase differences by detecting consistent non-zero phase lags, thereby reducing the contribution of zero-lag synchronization that may result from shared sources or artifacts [53]. The weighted phase lag index (wPLI) is an extension of the PLI that further improves sensitivity by weighting the contribution of phase differences based on their magnitude. This approach enhances the reliability of connectivity estimates, especially in scenarios where signals are weak or contaminated by noise, which is a common issue in SEEG recordings from deep brain structures [54, 53]. In our study, wPLI was primarily used for the resting-state data and the within-region connectivity analyses because the resting-state signals and the local interactions are more susceptible to confounding effects from volume conduction and common noise sources[53].

#### 5.7.3 Phase-slope-index analysis

To determine the direction of information flow across brain regions, we also used the phase-slope index (PSI) method to calculate the phase coherency slope between signals over a frequency band[50, 51]. The PSI offers a normalized measure of both the direction and strength of information flow between two signals. A positive PSI value suggests that the first signal drives the second, whereas a negative value indicates that the second signal drives the first. The magnitude of the PSI indicates the strength of this directed connectivity, with values ranging from −1 to 1. The PSI between two ROIs was computed for all inter-ROI electrode pairs for each trial, and then averaged over each experimental condition. We further conducted one sample t-tests against 0 for each condition to assess where there was significant information flow between the two brain regions. We also used the paired sample t-test to access possible differences between different conditions.

### 5.8 Inter-Subject Representational Similarity Analysis (IS-RSA)

To investigate whether the inter-subject similarity of functional connectivity patterns during resting-state was more comparable to different task state, we performed a Inter-Subject Representational Similarity Analysis (IS-RSA). This method allowed us to quantify the similarity between the brain’s functional architecture during task performance and its intrinsic architecture during the resting state[55, 56]. For each condition, we first organized functional connectivity data for each subject into a one-dimensional feature vector, where each value was assigned a sequential index based on its position within that subject’s ordered data sequence. For task conditions, this vector consisted of coherence values for each pair of brain regions. For the resting state, we used wPLI values. The Pearson correlation coefficient was calculated between the feature vectors of every pair of subjects to form the RSM. To ensure the robustness of our findings, we employed a bootstrap procedure (10,000 iterations), resampling the RDM elements with replacement to estimate the stability and confidence intervals of the observed correlations and their differences. We then calculated the Spearman’s rank correlation between the resting-state RSM vector and the task-state vectors. A non-parametric bootstrapping test (10,000 iterations) was implemented to assess the significance of the difference (Δ*ρ* = *ρ_intentional,res_ − ρ_lessintentional,res_*). A larger positive value for Δ*ρ* indicates a greater divergence of the circuit’s functional pattern from its intrinsic resting mode when processing intentional harm compared to less-intentional harm. It serves as a direct measure of intent-driven network reconfiguration. By repeatedly resampling the subject-pairs with replacement, we generated a distribution of correlation differences. The 95% confidence interval of this distribution was calculated. A statistically significant difference between the two correlations was concluded if this confidence interval did not span zero.

Also, to formally test our hypothesis that this network reconfiguration effect was stronger in adults than in children, we performed a non-parametric difference-of-differences permutation test. The null hypothesis for this test is that the magnitude of reconfiguration (Δ*ρ*) is the same across both age groups. We first calculated the observed difference between the reconfiguration scores of the two groups: *D_observed_*= Δ*ρ_adults_ −* Δ*ρ_children_*. We then constructed an empirical null distribution. All participants were pooled into a single group, and we performed 10,000 permutations. In each permutation, we randomly shuffled the group labels (‘adult’ or ‘child’) and recalculated the difference-of-differences (*D_permuted_*) using these shuffled labels. The p-value was calculated as the proportion of permutations where the absolute value of the permuted difference was greater than or equal to the absolute value of the observed difference (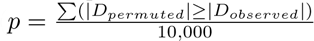).

### 5.9 Quantitative and statistical analysis

The statistical analysis of SEEG data was performed in Python (3.9.18) and the behavioral statistical test was performed in R (4.2.3). Independent-sample t-tests were used to assess the statistical significance between the behavior in different groups. Paired t-tests were also used to estimate the statistical significance between the BA or BGA and the baseline. Spearman correlation was used to calculate the rank correlation between patients’ age and the BGA as well as connectivity metrics. In the analysis of BGA signals, cluster-based permutation tests were used to calculate statistics corrected for multiple comparisons using permutations and cluster-level correction.

## Acknowledgement

This work was mainly supported by the Science and Technology Development Fund (FDCT) of Macau [0127/2020/A3, 0041/2022/A, 0112/2024/RIA2], the Natural Science Foundation of Guangdong Province(2021A1515012509), Shenzhen-Hong Kong-Macao Science and Technology Innovation Project (Category C) (SGDX2020110309280100), and the MYRG of University of Macau (MYRG2022-00188-ICI). We also thank all research assistants who provided general support in data collection.

## Data and Code Availability

The SEEG data used in this study cannot be made publicly available due to privacy protection of the patients, but can be requested from the corresponding author [Haiyan Wu]. Code for the present data analyses is available at the GitHub repository: https://github.com/andlab-um/TPP_sEEG.

