## Supplemental Materials for "Intracranial Insights into the Developing Neural Basis of Moral Punishment"

### Supplementary Materials

July 22, 2025

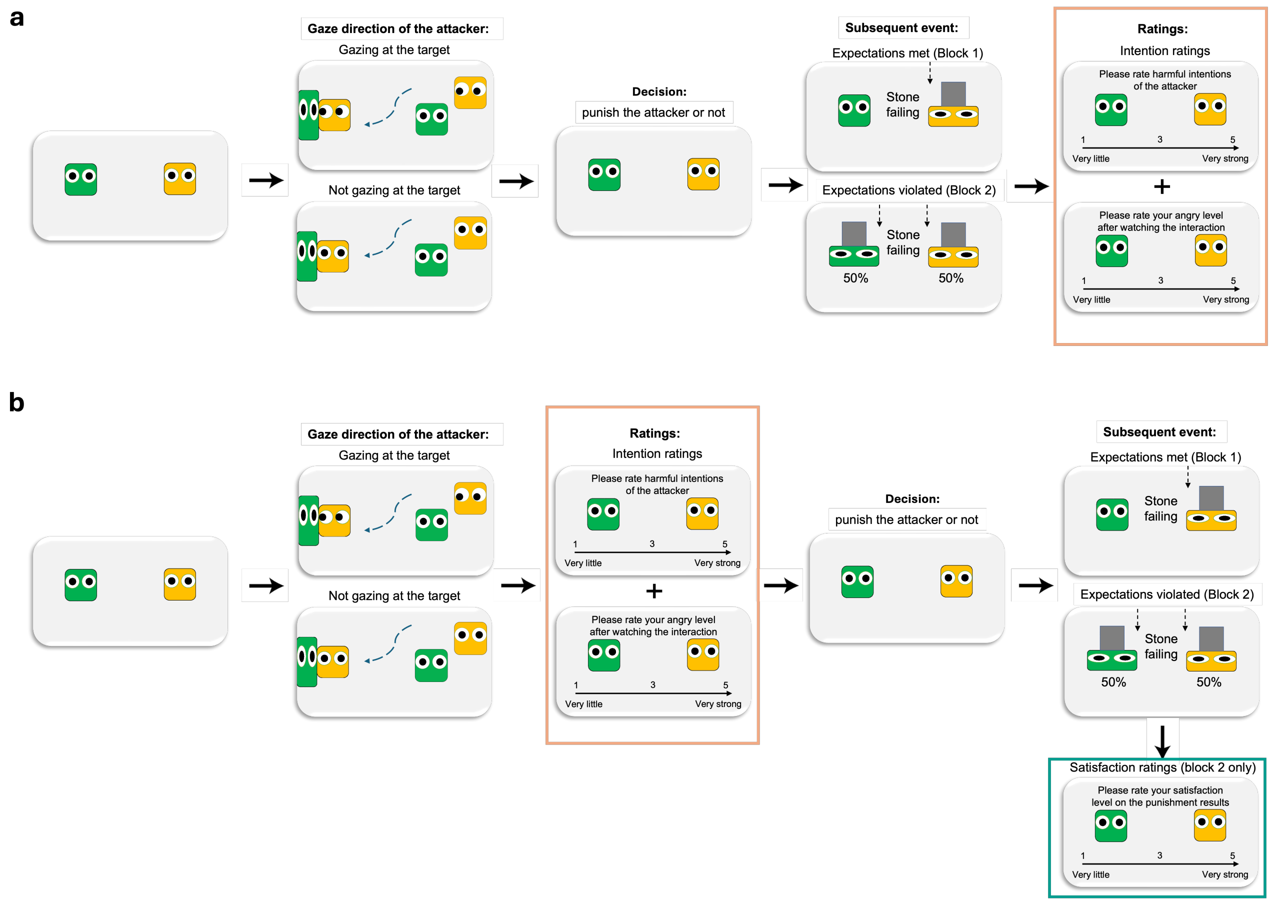


Figure S1: **The experimental paradigm used in behavioral Exp. 1 and Exp. 2. (a)** The behavioral paradigm of Exp.1. Participants were asked to rate the perceived harmful intention and their angry level after completing the whole task. **(b)** The behavioral paradigm of Exp.2. In Exp. 2 participants were asked to do both ratings right after the interaction video clip in each trial, before decision-making. There was also an extra rating on the satisfaction level of the punishment results in block 2 of Exp.2.

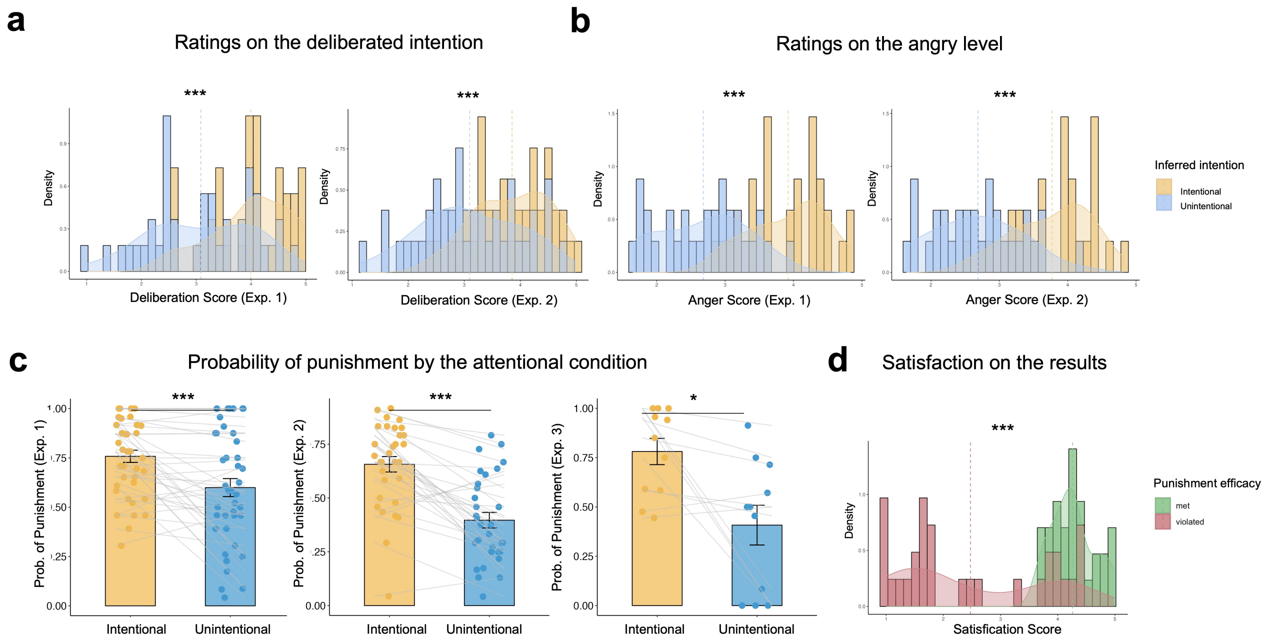


Figure S2: **The behavioral validation of Exp. 1 and Exp. 2 suggest that the paradigm can achieve the manipulation of the attacker's intention and punishment efficacy. * denoted significant one sample t test ($p < 0.05$) against 0, ** denoted test results $p < 0.01$, and *** denoted test results $p < 0.001$, same below.** **(a)** The ratings of the deliberated intention of the attackers on the interaction. The ratings were higher in the intentional condition in both Exp.1 and Exp.2. **(b)** The ratings of participants' anger levels after watching the video. The ratings were higher in the intentional condition in both Exp.1 and Exp.2. **(c)** The probability of punishment decision. Participants tended to punish more when the attacker was paying attention to the target in both Exp. 1, Exp. 2 and Exp. 3. **(d)** The effects of expectation violation of the results on participants' decision in the next trial. **(e)** The satisfaction level of the punishment results (Exp.2). Participants were more satisfied when the punishment results were consistent with their expectations.

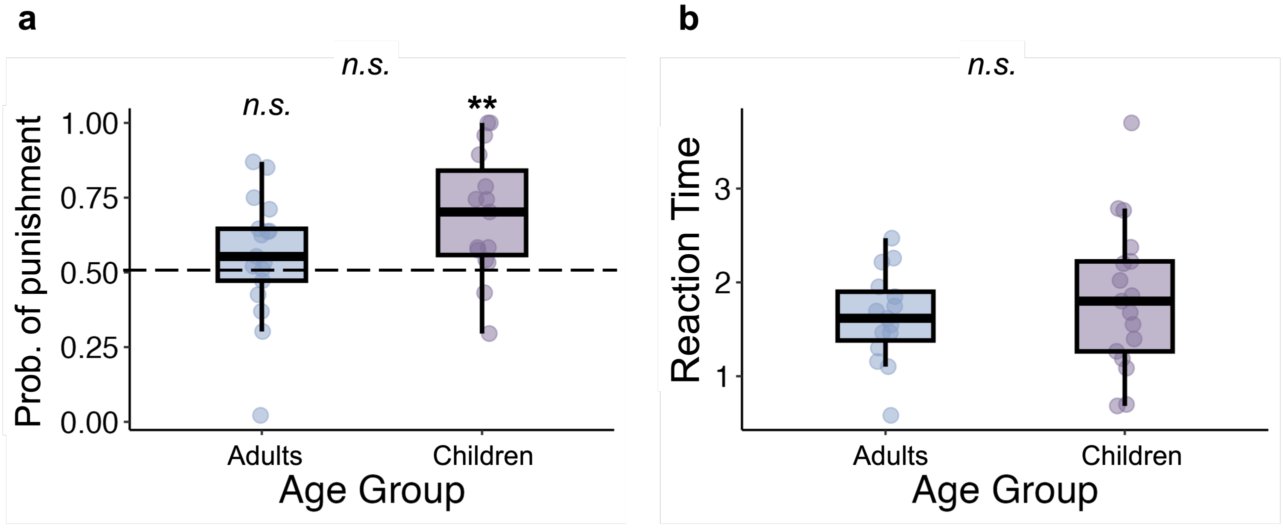


Figure S3: **The probability of punishment and reaction time of the SEEG patients in adult and children group.**

| Group | Regressor | Mean estimate | t value | p value | 95 % CI |
|---|---|---|---|---|---|
| **Healthy Adults (N = 71)** | Trial | 0.00 | -1.52 | 0.132 | [-0.003, 0.000] |
| | Block | -0.05 | -2.08 | 0.040* | [-0.10, -0.002] |
| **Healthy Children (N = 11)** | Trial | 0.00 | 0.85 | 0.412 | [-0.005, 0.01] |
| | Block | 0.12 | 1.60 | 0.140 | [-0.05, 0.29] |
| **SEEG Adults (N = 16)** | Trial | 0.00 | 0.05 | 0.96 | [-0.01, 0.01] |
| | Block | -0.01 | -0.24 | 0.82 | [-0.13, 0.11] |
| **SEEG Children (N = 13)** | Trial | 0.00 | 0.18 | 0.86 | [-0.05, 0.06] |
| | Block | 2.02 | 1.32 | 0.25 | [-1.67, 5.71] |

Table S1: **Model estimates of M0.** Linear regression model were used to predict patients' choice. This model only contained two predictors (trial number and block). Results showed that neither of the two factors affected decision making, which eliminated the possible learning effect.

| Group | Regressor | Mean estimate | t value | p value | 95 % CI |
|---|---|---|---|---|---|
| **Healthy Adult** | Intention | 0.19 | 6.11 | 0.000*** | [0.13, 0.25] |
| | Punishment efficacy | -0.03 | -2.83 | 0.006* | [-0.06, -0.01] |
| **Healthy Children** | Intention | 0.39 | 2.83 | 0.02* | [0.08, 0.68] |
| | Punishment efficacy | 0.008 | 0.24 | 0.81 | [-0.07, 0.09] |
| **SEEG Adult** | Intention | 0.09 | 2.27 | 0.037* | [0.006, 0.17] |
| | Punishment efficacy | -0.09 | -2.50 | 0.024* | [-0.16, -0.01] |
| **SEEG Children** | Intention | 0.01 | 0.33 | 0.747 | [-0.08, 0.11] |
| | Punishment efficacy | -0.06 | -1.17 | 0.267 | [-0.17, 0.05] |

Table S2: **Model estimates of M1.** Linear regression were used to predict TPP decisions with two factors (attacking intention and punishment efficacy) with the reaction times (RTs), trials, and blocks controlled. The attacking intention was coded as whether the attacker was paying attention (=1) or not paying attention (=0) to the target. The punishment efficacy was coded as whether the punishment results met (=1) or violated (=0) their expectation of punishing the attacker. This model also included RT and block number as controlling variables. The inferred attacking intention can predict the decision in both adult sample and the healthy children sample but not in SEEG children sample. Further, we also found that the punishment efficacy can only predict the decision in adults but not in children.

| Group | Regressor | Mean estimate | t value | p value | 95 % CI |
|---|---|---|---|---|---|
| Healthy Adults | Attention | 0.147 | 3.794 | 0.000* | [0.070, 0.224] |
| | RT | -0.001 | -0.087 | 0.931 | [-0.033, 0.030] |
| | Punishment Efficacy | -0.025 | -1.231 | 0.223 | [-0.066, 0.016] |
| | Block | -0.078 | -1.737 | 0.087 | [-0.167, 0.012] |
| | Trial | 0.000 | 0.311 | 0.756 | [-0.002, 0.002] |
| | Punishment Efficacy:Block | -0.023 | -0.547 | 0.586 | [-0.105, 0.060] |
| | Attention:Punishment Efficacy | 0.012 | 0.399 | 0.691 | [-0.049, 0.074] |
| | Attention:Punishment Efficacy:Block | -0.056 | -0.842 | 0.403 | [-0.191, 0.078] |
| Healthy Children | Attention | 0.424 | 3.277 | 0.008* | [0.136, 0.712] |
| | RT | -0.068 | -1.518 | 0.160 | [-0.168, 0.032] |
| | Punishment Efficacy | -0.029 | -0.592 | 0.567 | [-0.137, 0.079] |
| | Block | 0.025 | 0.182 | 0.859 | [-0.281, 0.331] |
| | Trial | -0.001 | -0.285 | 0.782 | [-0.010, 0.008] |
| | Punishment Efficacy:Block | 0.137 | 1.268 | 0.237 | [-0.108, 0.382] |
| | Attention:Punishment Efficacy | 0.004 | 0.096 | 0.925 | [-0.099, 0.108] |
| | Attention:Punishment Efficacy:Block | -0.112 | -0.831 | 0.433 | [-0.431, 0.207] |
| SEEG Adults | Attention | 0.102 | 1.984 | 0.065 | [-0.007, 0.212] |
| | RT | -0.014 | -0.475 | 0.641 | [-0.078, 0.049] |
| | Punishment Efficacy | 0.038 | 0.528 | 0.605 | [-0.114, 0.189] |
| | Block | 0.092 | 0.975 | 0.344 | [-0.108, 0.291] |
| | Trial | 0.001 | 0.502 | 0.622 | [-0.005, 0.008] |
| | Punishment Efficacy:Block | -0.206 | -1.908 | 0.076 | [-0.437, 0.024] |
| | Attention:Punishment Efficacy | -0.193 | -2.028 | 0.061 | [-0.396, 0.010] |
| | Attention:Punishment Efficacy:Block | 0.198 | 1.232 | 0.237 | [-0.144, 0.539] |
| SEEG Children | Attention | -0.060 | -1.328 | 0.207 | [-0.158, 0.038] |
| | RT | 0.020 | 0.893 | 0.388 | [-0.028, 0.068] |
| | Punishment Efficacy | 0.052 | 1.098 | 0.292 | [-0.050, 0.154] |
| | Block | 0.073 | 0.656 | 0.523 | [-0.167, 0.313] |
| | Trial | 0.002 | 0.901 | 0.384 | [-0.003, 0.008] |
| | Punishment Efficacy:Block | -0.159 | -1.465 | 0.167 | [-0.394, 0.076] |
| | Attention:Punishment Efficacy | -0.018 | -0.236 | 0.817 | [-0.184, 0.147] |
| | Attention:Punishment Efficacy:Block | -0.036 | -0.172 | 0.868 | [-0.514, 0.443] |

Table S3: **Model estimates of M2.** Linear regression were used to predict TPP decisions with two factors (attacking intention and punishment efficacy) and their interaction term with the reaction times (RTs), trials, and blocks controlled. The attacking intention was coded as whether the attacker was paying attention (=1) or not paying attention (=0) to the target. The punishment efficacy was coded as whether the punishment results met (=1) or violated (=0) their expectation of punishing the attacker. This model also included RT and block number as controlling variables. No significant interaction were found on those factors.

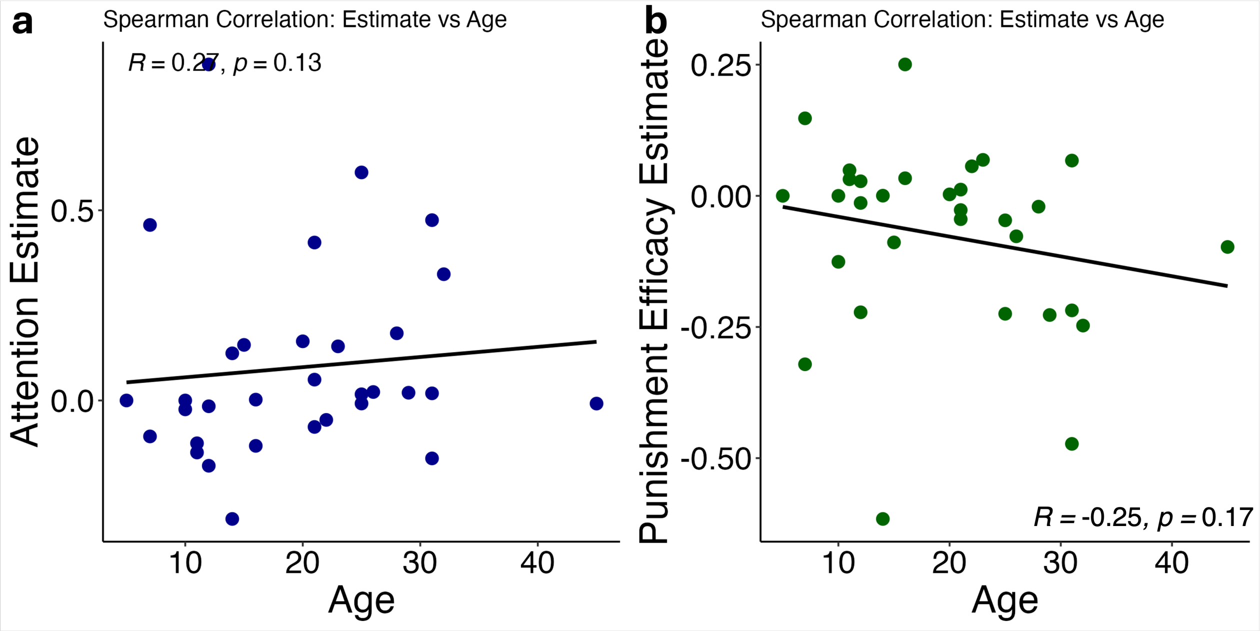


Figure S4: **Correlation pattern of behavioral model estimate and patients age.** **(a)** Correlation between intention estimates and age. **(b)** Correlation between punishment efficacy estimates and age.

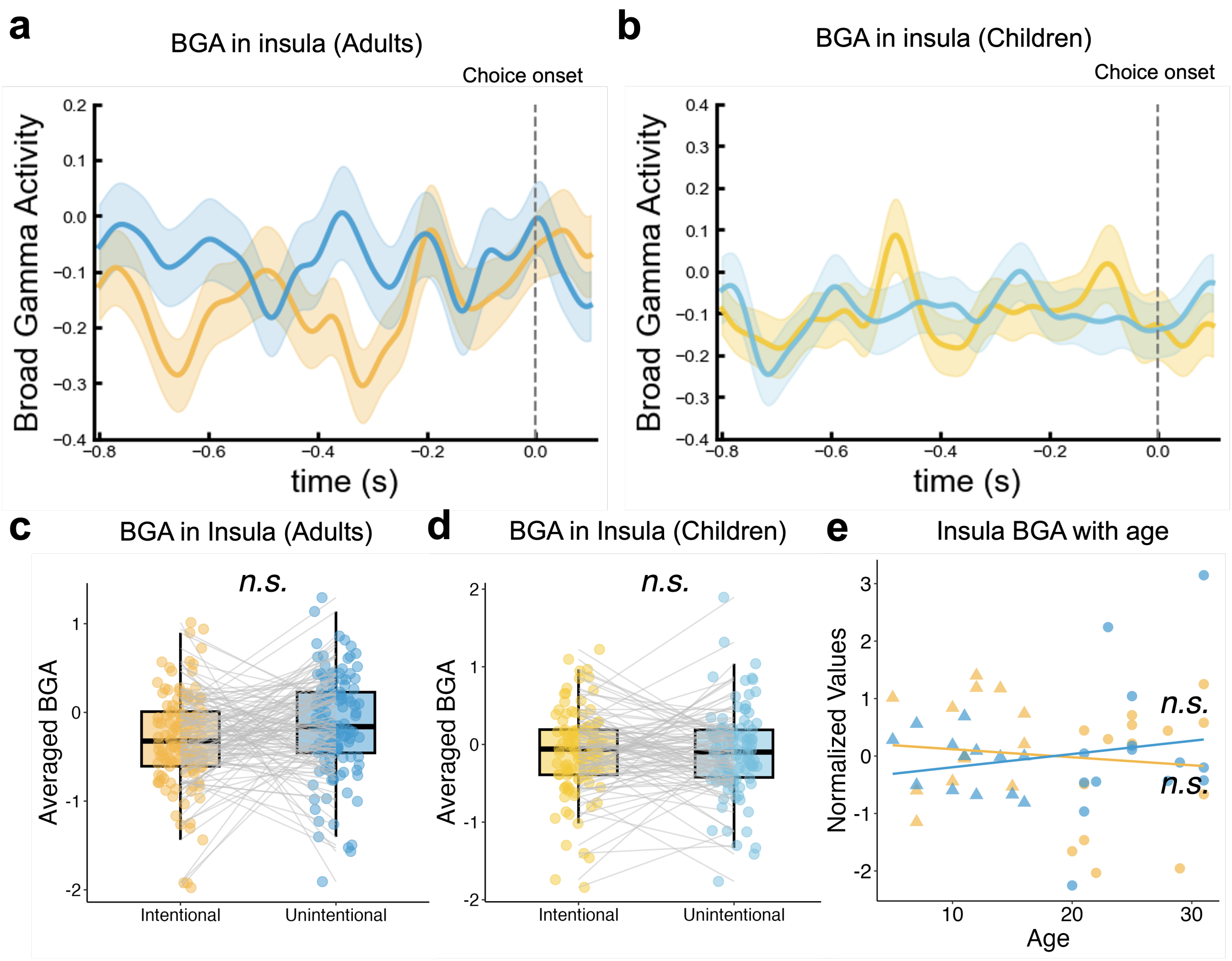


Figure S5: **No significant difference in the BGA of insula before the choice onset in different condition of the attacker's intention.** **(a)** The neural activity in the insula before choice onset in the adults group. The yellow and blue lines indicate neural activity during decision-making in the intentional and less intentional conditions, respectively. The gray shades in the background indicate the period when there was a significant difference in neural activity between the two conditions. Error ribbons represent the standard error of the mean. The zero point of the time series is the moment when the patients make the punishment decision. **(b)** The neural activity in the insula before choice onset in the children group. **(c)-(d)** The averaged BGA in the insula before choice onset in the adult and children group. **(e)** correlation between insula BGA and age. Triangles represented the averaged BGA power for child patients during decision.

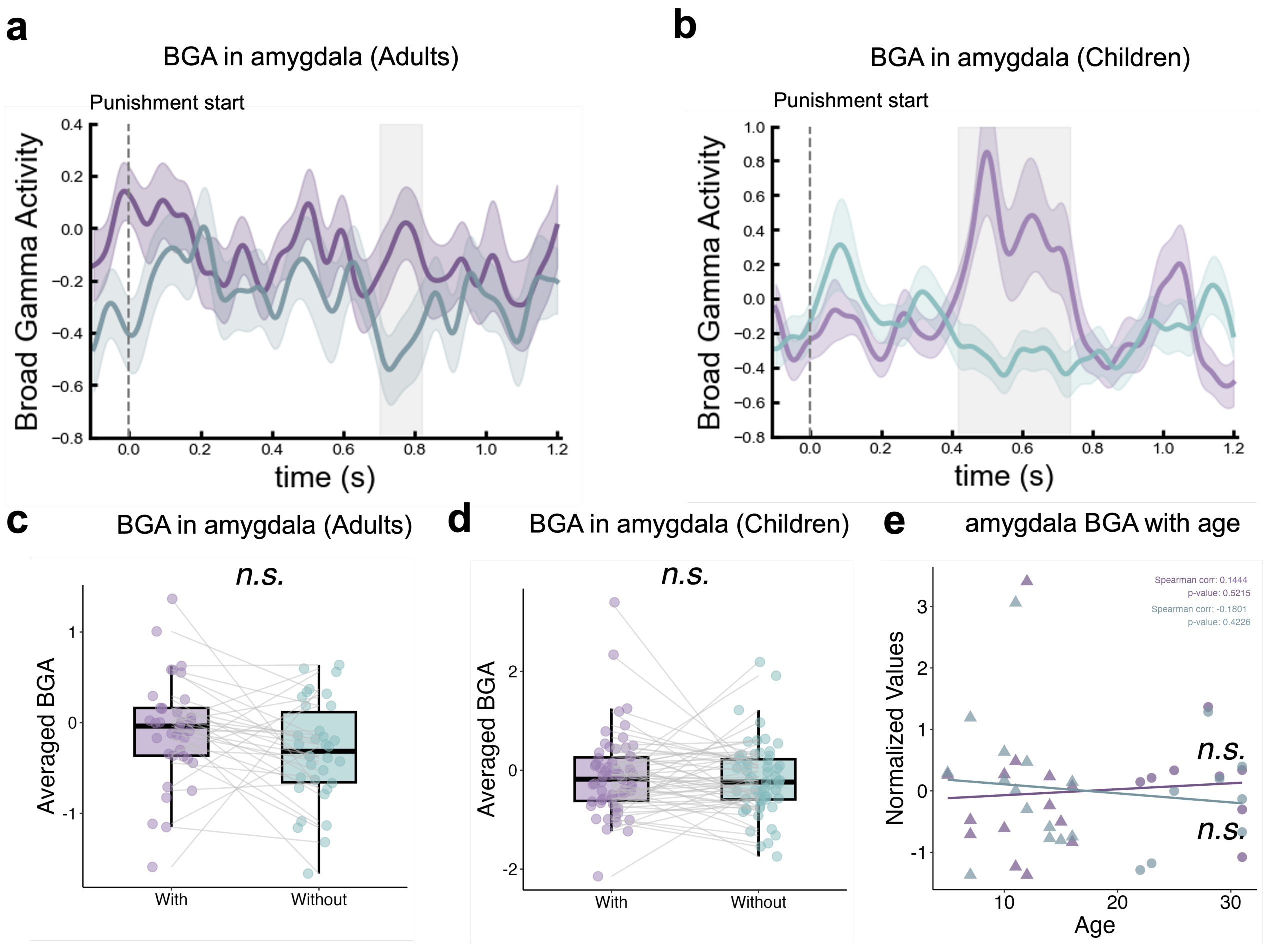


Figure S6: **No significant difference in the BGA of amygdala during punishment result presentation.** **(a)** The neural activity in the amygdala after result presentation in the adults group, represented by purple (with efficacy) and gray (without efficacy) lines. Gray-shaded areas indicate periods of significant differences in activity between the conditions. Error bars represent the standard error of the mean. The time series is aligned to the moment of the punishment result presentation (time zero). **(b)** Neural activity in the amygdala after the presentation of the results in the children group. **(c)-(d)** The averaged BGA in the insula and vmPFC after the presentation of the results in the adult and children group. **(e)** correlation between amygdala BGA and age. Triangles represented the averaged BGA power for child patients during decision.

| SBJ | vmpfc*amygdala | vmpfc*insula | insula*amygdala | vmpfc*ipl |
|---|---|---|---|---|
| A01 | / | / | 14 | / |
| A02 | / | / | 8 | / |
| A03 | 18 | 27 | 54 | 51 |
| A04 | 75 | 105 | 35 | / |
| A05 | 108 | 198 | 66 | 306 |
| A06 | / | 370 | / | 296 |
| A07 | 60 | 240 | 36 | 160 |
| A08 | / | / | / | / |
| A09 | 21 | 196 | 84 | / |
| A10 | 10 | 30 | 12 | 170 |
| A11 | / | 102 | / | 374 |
| A12 | / | 18 | / | 189 |
| A13 | / | / | / | / |
| A14 | 24 | 18 | 12 | 36 |
| Total Adult | 316 | 1304 | 321 | 1582 |
| C01 | / | / | 33 | / |
| C02 | 44 | 220 | 20 | / |
| C03 | / | / | 10 | / |
| C04 | 18 | 42 | 21 | 72 |
| C05 | 12 | 68 | 51 | 168 |
| C06 | / | / | 55 | / |
| C07 | / | / | / | / |
| C08 | / | / | 28 | / |
| C09 | 80 | 144 | 45 | 336 |
| C10 | 28 | 77 | 44 | / |
| C11 | 40 | 50 | 80 | 75 |
| C12 | 20 | 50 | 40 | 70 |
| C13 | 250 | 100 | 10 | / |
| C14 | / | / | 16 | / |
| Total Child | 492 | 751 | 453 | 721 |

Table S4: **Electrode pairs in each patient.** To further investigate functional connectivity across brain regions, we computed the phase-locking value (PLV), coherence, and phase synchrony index (PSI) between two regions of interest (ROIs) for all inter-ROI electrode pairs within each condition.

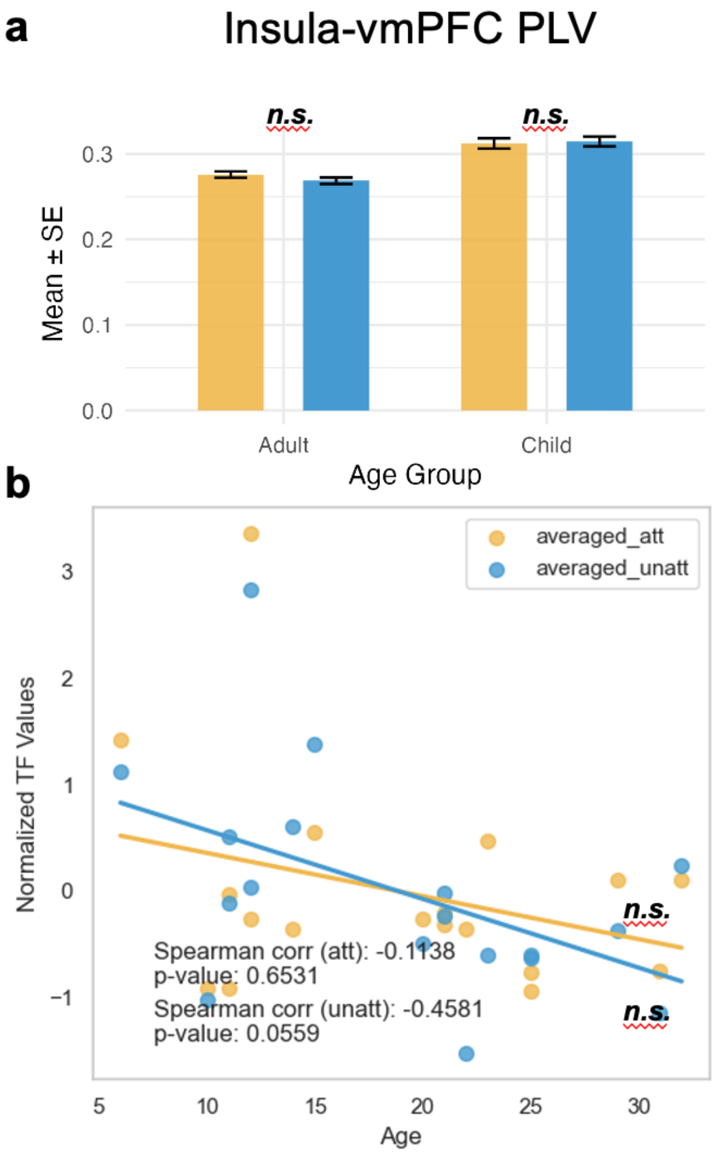


Figure S7: **The insula-vmPFC PLV. (a)** The PLV between insula and vmPFC during decision-making in different age groups' intentional and less intentional conditions. * denoted significant one sample t test ($p < 0.05$) against 0, ** denoted test results $p < 0.01$, and *** denoted test results $p < 0.001$, same below. **(c)** The Spearman correlation between the insula-vmPFC PLV and patients' age for the intentional and less intentional conditions of the attacker's intention.

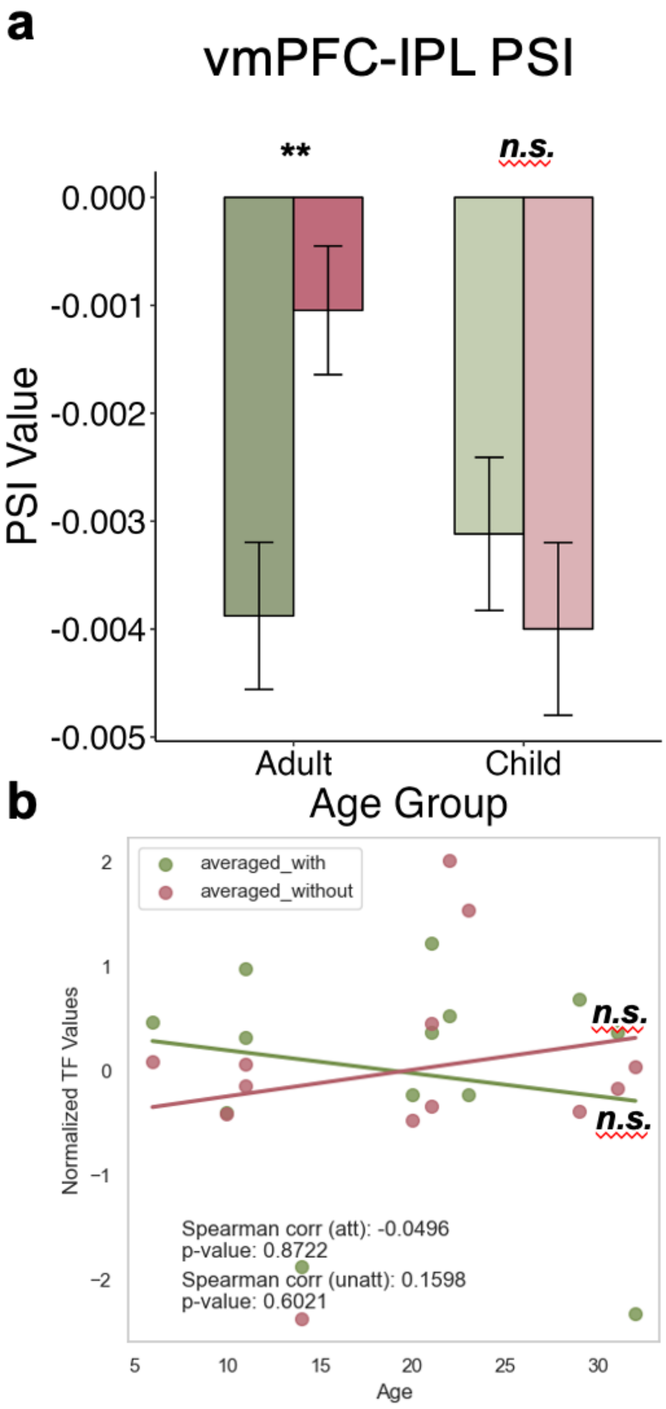


Figure S8: **The vmPFC-IPL PSI values. (a)** The PSI value between vmPFC and IPL during decision-making in different age groups' with efficacy and without efficacy conditions. * denoted significant one sample t test ($p < 0.05$) against 0, ** denoted test results $p < 0.01$, and *** denoted test results $p < 0.001$, same below. **(b)** The Spearman correlation between the vmPFC-IPL PSI and patients' age for the with efficacy and without efficacy of the attacker's intention.

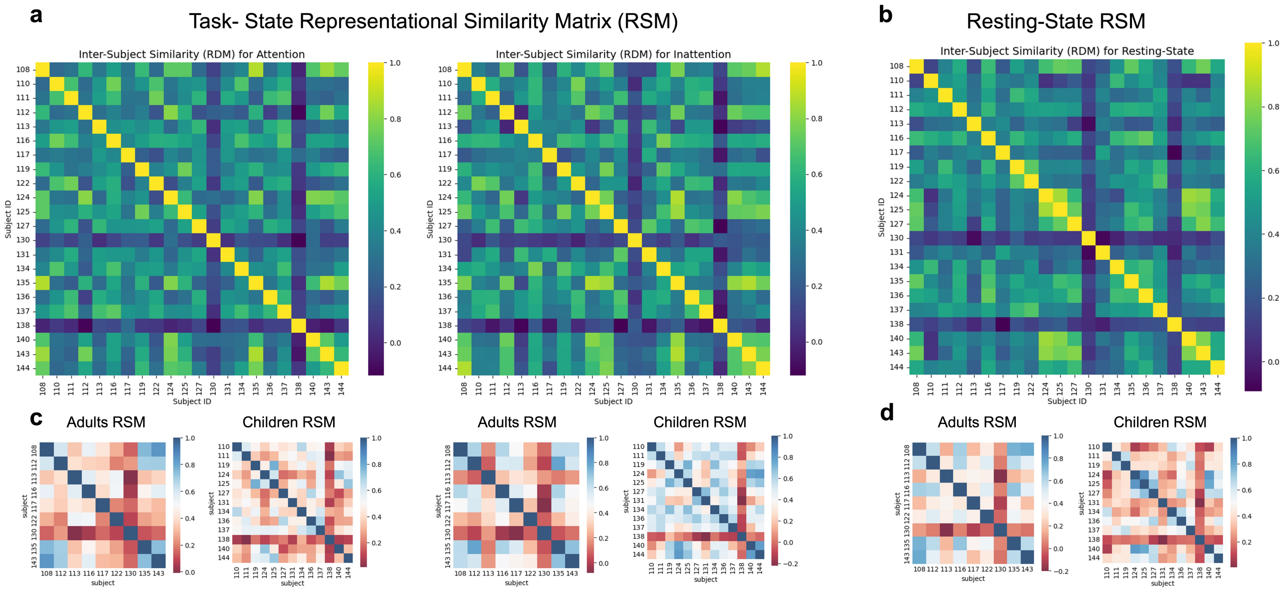


Figure S9: **Intersubject Representational Similarity Matrices (RSMs) for Amygdala–Insula Connectivity. (a) & (b)** These figures displays the first-order Representational Similarity Matrices (RSMs) used in the main analysis to compare task-evoked and resting-state neural patterns. A single cell $(i, j)$ in each matrix represents the Pearson correlation $(r)$ between the vectorized amygdala–insula connectivity patterns of subject $i$ and subject $j$. **(c) & (d)** Each matrix shows the intersubject similarity structure for a specific condition and age group. The matrices are shown for adults (top row) and children (bottom row) across the three conditions: Intentional harm, Less-intentional harm, and Resting state. Warmer colors (red) indicate higher intersubject similarity (i.e., subjects exhibited more similar neural patterns), while cooler colors (blue) indicate lower similarity or dissimilarity. These RSMs formed the basis for the second-order similarity analysis and the subsequent permutation testing reported in the main text.
